# MF-PENS: an efficient and versatile nuclear extraction method for plant single-cell sequencing

**DOI:** 10.1101/2025.03.14.643295

**Authors:** Lin Du, Qing Liu, Hongling Zhou, Mengyue Zhao, Fuyan Liu, Cheng Tong, Mengjiao Zhang, Hao Yu, Zhe Liang, Jingmin Kang

## Abstract

The widespread adoption of single-cell transcriptomics in plants remains constrained by the challenges of isolating high-quality nuclei from tissues with rigid cell walls and complex metabolomes. Here, we present **m**ethanol **f**ixation and **P**ercoll-**e**nriched **n**uclei **s**orting (MF-PENS), a robust strategy designed to tackle these limitations. Distinct from conventional protocols, MF-PENS circumvents the need for fluorescence activated cell sorting (FACS) and effectively reduce RNA leakage, consistently yielding high-quality single-nucleus RNA sequencing (snRNA-seq) data. MF-PENS has been validated across various plant tissues, including fresh and frozen rice roots and leaves, as well as recalcitrant samples such as soybean root nodules, soybean seeds, and tomato fruits. Benchmarking results demonstrate that MF-PENS significantly enhances gene capture efficiency, cell clustering performance, and cell type annotation accuracy. With its broad applicability and superior performance, MF-PENS represents a robust and versatile tool for plant single-cell research, facilitating deeper insights into plant development and biology processes.

## Introduction

Cells constitute the fundamental units of growth and development in multicellular organisms. The formation of complex tissues and organs is driven by the dynamic interactions among diverse cell types and their specialized functions.^1^ Recent studies have highlighted that even morphologically similar cells within the same tissue can exhibit substantial heterogeneity, characterized by distinct gene expression signatures.^2,3^ Single-cell RNA sequencing (scRNA-seq) technologies have revolutionized genomic and transcriptomic profiling by offering single-cell resolution. Unlike traditional bulk sequencing, scRNA-seq enables the detection of rare but biologically significant cell populations, the dissection of heterogeneity among phenotypically indistinguishable cells, and the exploration of stochastic variation within genetically identical populations.^4^ Furthermore, these technologies facilitate precise cell type annotation, functional characterization, and the inference of cellular dynamics. Consequently, scRNA-seq has been widely adopted across diverse disciplines, including cancer biology, immunology, stem cell research, neuroscience, infectious diseases, and plant stress biology.^5-9^

In plant biology, pioneering scRNA-seq applications focused on establishing foundational resources, such as the construction of high-resolution cellular atlases. For instance, Liu *et al.* and Zhang *et al.* established comprehensive single-cell atlases of rice root tips.^10,11^ More recently, the field has pivoted toward elucidating the molecular mechanisms governing plant responses to environmental stresses. Liu *et al.* utilized single-cell transcriptomics to dissect the response of *Arabidopsis thaliana* to osmotic stress,^9^ while Li *et al.* characterized the cellular responses to salt stress in cotton. Additionally, Swift *et al.* leveraged these tools to unravel key mechanisms driving the evolution of efficient photosynthetic pathways in C4 plants.^12^ These studies represent a paradigm shift in plant single-cell research—transitioning from cellular atlas construction to the investigation of biological functions and regulatory mechanisms. However, the application of single-cell technologies in plants continues to lag behind their widespread adoption in animal models.^13^ A primary impediment lies in the intrinsic structural and biochemical complexity of plant cells^14,15^: they are encased in rigid cell walls and often contain high concentrations of interfering complex secondary metabolites, such as polysaccharides and polyphenols. These physicochemical features impose substantial technical challenges for isolating high-quality protoplasts or nuclei from plant tissues, which are essential for scRNA-seq or single-nucleus RNA sequencing (snRNA-seq).^16-19^

Protoplast preparation typically entails prolonged enzymatic digestion to degrade the cell wall matrix.^20-22^ The efficiency of this process is highly dependent on species and tissue characteristics, precluding the establishment of universal, standardized protocols. Crucially, cells remain metabolically active during enzymatic digestion; extended exposure to enzymatic stress can induce transcriptional artifacts—such as the ectopic activation of stress-response genes—preferential depletion of rare or enzyme-sensitive cell populations. These confounding factors ultimately compromise fidelity and completeness of the cellular landscape reconstructed by scRNA-seq. Furthermore, protoplast isolation strictly mandates the use of fresh tissue, imposing strict temporal constraints on sample processing. This requirement limits experimental flexibility and prevents the retrospective analysis of archived or frozen specimens.

To circumvent the limitations inherent to protoplast-based methods, snRNA-seq has emerged as a promising alternative. Farmer *et al.* demonstrated the reliability of snRNA-seq in plant research using *Arabidopsis* root tissues, showing its capacity to resolve cell subtypes often obscured in protoplast-derived datasets.^23^ However, the isolation of high-quality plant nuclei remains a significant technical bottleneck. Current approaches, such as those applied to *Arabidopsis* and wheat roots,^23,24^ typically involve crude nuclear extraction followed by purification using fluorescence activated cell sorting (FACS), which enables snRNA-seq. Despite their effectiveness, these methods depend on specialized instrumentation (flow cytometer) that is not universally accessible, thereby imposing a high barrier to entry for many laboratories^12,25,26^. Although some studies have attempted to perform snRNA-seq without FACS, they frequently struggle to maintain the integrity of nucleus.^27^ Physical damage during extraction often results in RNA leakage, which manifests as low library complexity (few genes detected per nucleus) or the loss of nuclei during quality control due to insufficient transcript capture^28-31^. Notably, although a few studies have employed methanol fixation in snRNA-seq, their primary purpose was solely sample storage, with no indication that this treatment enhances data quality; furthermore, these methods still rely on FACS for nuclear purification. Moreover, methanol fixation protocols have not clearly specified critical parameters (such as fixation time and methanol concentration) or standardized operating procedures, limiting reproducibility and broad applicability.^32^ Collectively, these technical hurdles continue to impede the widespread adoption of single-cell omics in plant research.

To address this gap, we report a robust and versatile nuclei isolation strategy named MF-PENS that integrates methanol fixation with density gradient centrifugation. Designed to circumvent existing technical barriers, this method provides a comprehensive solution featuring (1) full compatibility with frozen tissues, (2) a FACS–free purification workflow, and (3) a fixation-based preservation system that ensures optimal nuclear integrity and RNA stability. We demonstrate the universality of this approach across a diverse array of plant tissues—including roots, leaves, root nodules, fruits, seeds, and callus—consistently yielding high-quality input for snRNA-seq. By establishing a user-friendly, cost-effective, and highly reproducible standard, this method empowers the research community to fully leverage single-cell resolution for advancing plant developmental biology, functional genomics, and agricultural innovation.

## Results

### Establishment of the MF-PENS workflow for plant single-nucleus sequencing

To optimize a nuclei isolation protocol suitable for high-quality plant snRNA-seq, we systematically benchmarked four distinct fixation conditions combined with the Percoll-enhanced nuclei separation (PENS) purification: 0.1% formaldehyde fixation (0.1%FF-PENS), 1% formaldehyde fixation (1%FF-PENS), 100% methanol fixation (MF-PENS), and a no-fixation control (NT-PENS). In detail, plant tissues were sectioned into approximately 0.5 cm^²^ fragments and subjected to their respective fixation treatment, with the exception of the NT-PENS method which was proceeded directly to the next step without fixation. Subsequently, nuclei were released and purified via Percoll density gradient centrifugation. The isolated nuclei then underwent microfluidic-based barcoding, library construction, and high-throughput sequencing **(Figure 1A)**. We evaluated the performance of these methods by accessing data quality metrics and downstream analytical utility, including clustering resolution and the identification of differentially expressed genes (DEGs). Furthermore, we validated the robustness and versatility of MF-PENS across a diverse array of sample types, ranging from standard vegetative tissues (roots and leaves) to recalcitrant reproductive and symbiotic tissues, including root nodules, fruits, seeds, and callus **(Figure 1A)**.

**Figure 1.**
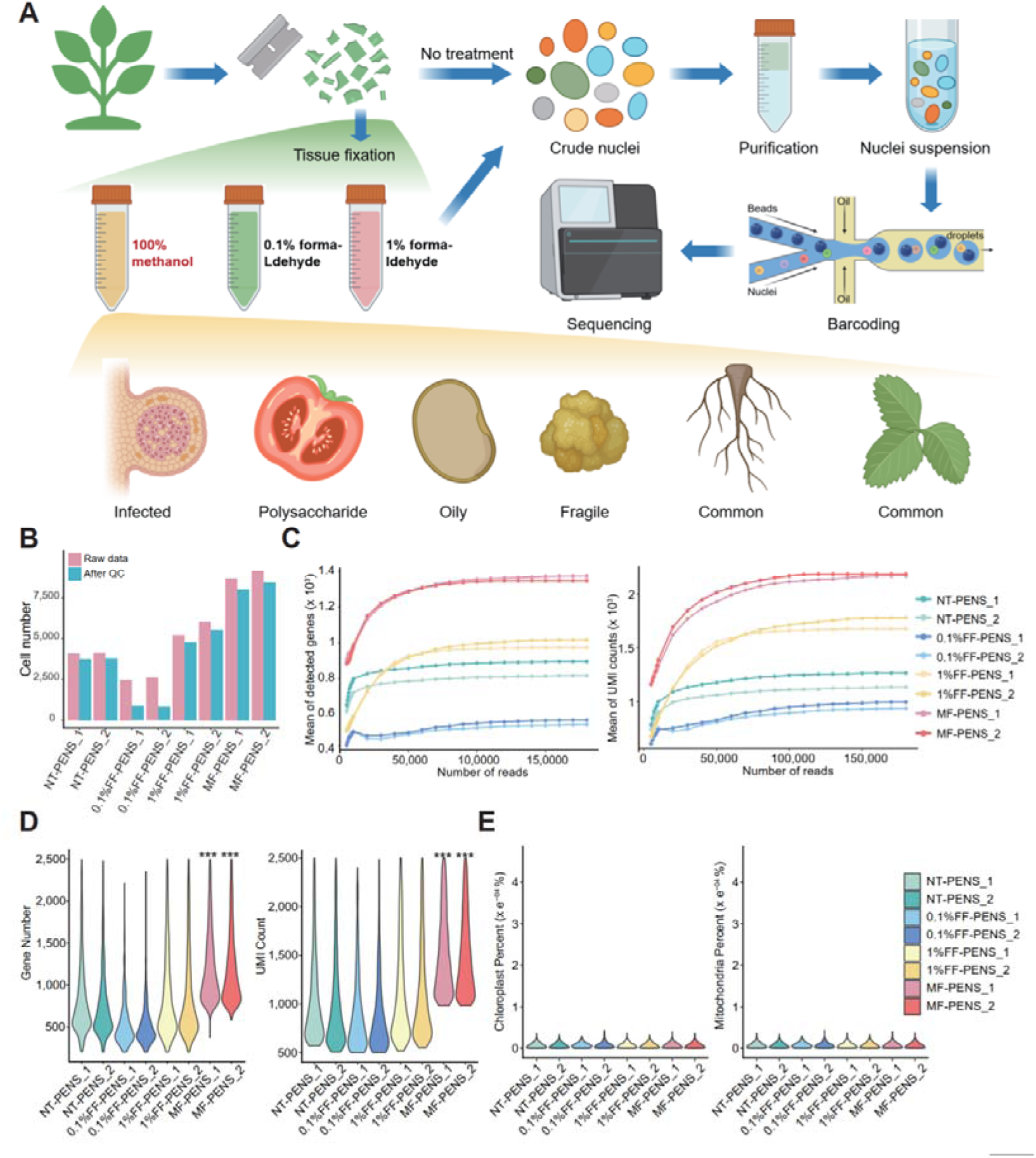
Design principle and performance evaluation of MF-PENS on rice roots. **(A)** Workflow diagrams for NT-PENS, 0.1%FF-PENS, 1%FF-PENS, and MF-PENS, illustrating distinct fixation and extraction strategies. The yellow shade indicates the application of MF-PENS in various plants, demonstrating its wide adaptability and robustness. **(B)** Cell counts of eight samples before (pink)/ after (blue) quality control, with two biological replicates for each method. **(C)** Stepwise increases in the number of detected genes and UMIs across down-sampled sequencing depths (sequencing depth vs. gene detection efficiency). Points indicate mean values at the corresponding depth, and colors represent snRNA-seq data generated by different methods. **(D)** Violin plots showing the distributions of UMIs and detected genes across four methods. *** indicates that MF-PENS yields significantly higher values (*P* < 0.05) compared with the other methods. **(E)** Mitochondrial and chloroplast percentages (y-axis values × 10^-4^%) for the eight biological samples corresponding to the four methods.

### MF-PENS yields high-quality single-nucleus transcriptomes

To assess the impact of fixation methods on snRNA-seq data quality, we systematically benchmarked four approaches in rice (*Oryza sativa*) root tissues (**Figure 1B-E, Supplementary Table S1**). All experiments were performed in biological replicates to ensure reproducibility. During data preprocessing, stringent quality control (QC) criteria were applied, including the removal of doublets/multiplets, empty droplets, and low-quality nuclei exhibiting high mitochondrial content (> 5%) or signs of enzymatic degradation. Notably, MF-PENS retained the highest number of high-quality nuclei (9,170 and 8,468) following identical QC procedures**(Figure 1B)**. Sequencing saturation analysis also demonstrated that MF-PENS consistently outperformed other methods across all examined read depths. MF-PENS replicates exhibited superior gene detection curves, recovering significantly more genes (*P* <0.05, Student’s *t*-test) and producing higher unique molecular identifier counts (UMIs) than other methods at equivalent depths. These findings demonstrate that MF-PENS confers enhanced transcript capture efficiency, establishing it as a robust approach for plant snRNA-seq **(Figure 1C)**.

Quantitative assessment confirmed that MF-PENS generated significantly higher post-QC UMIs and gene numbers compared to other methods (*P* <0.05), with high reproducibility **(Figures 1D and S1A)**. To assess nuclear purity, we quantified the proportions of chloroplast- and mitochondrial-derived transcripts **(Figure 1E)**. All methods exhibited low levels of organelle-derived transcripts, indicating minimal cytosolic contamination and high integrity of the extracted nuclear RNA. Additionally, Pearson correlation analysis **(Figure S1B)** demonstrated strong global similarity across the datasets. Interestingly, while 0.1%FF-PENS showed the highest correlation with NT-PENS, MF-PENS exhibited a distinct transcriptomic profile. We hypothesize that this divergence reflects the superior capacity of MF-PENS to preserve fragile, transcriptionally active nuclei, thereby capturing biologically relevant patterns underrepresented in other methods. Collectively, these results highlight the superiority of MF-PENS performance in generating optimal snRNA-seq data quality, characterized by higher nuclei yield, robust library complexity, and enhanced transcriptomic fidelity.

### MF-PENS enhance cell clustering performance and biological DEGs identification

While UMIs and gene detection rates are fundamental quality indicators, robust clustering and accurate marker identification are pivotal prerequisites for resolving biological heterogeneity.^33-36^ To evaluate clustering performance, we integrated biological replicates for each method. Uniform manifold approximation and projection dimensionality reduction (UMAP) visualization revealed negligible batch effects, validating experimental reproducibility and ensuring the rigor of downstream comparisons **(Figure 2A)**. At a fixed resolution yielding nine clusters, MF-PENS and 1%FF-PENS produced discrete, well-separated populations, whereas NT-PENS and 0.1%FF-PENS exhibited significant aggregation and boundary overlap, indicating compromised clustering capacity **(Figure 2B)**. This visual assessment was corroborated quantitatively by the Modularity score^37-39^, a graph-based metric that quantifies the contrast between intra- and inter-cluster connectivity, with higher values indicating better clustering result. MF-PENS achieved the highest Modularity score surpassing all other methods **(Figure 2B)**, thereby confirming its superior capacity to resolve distinct cell populations.

**Figure 2.**
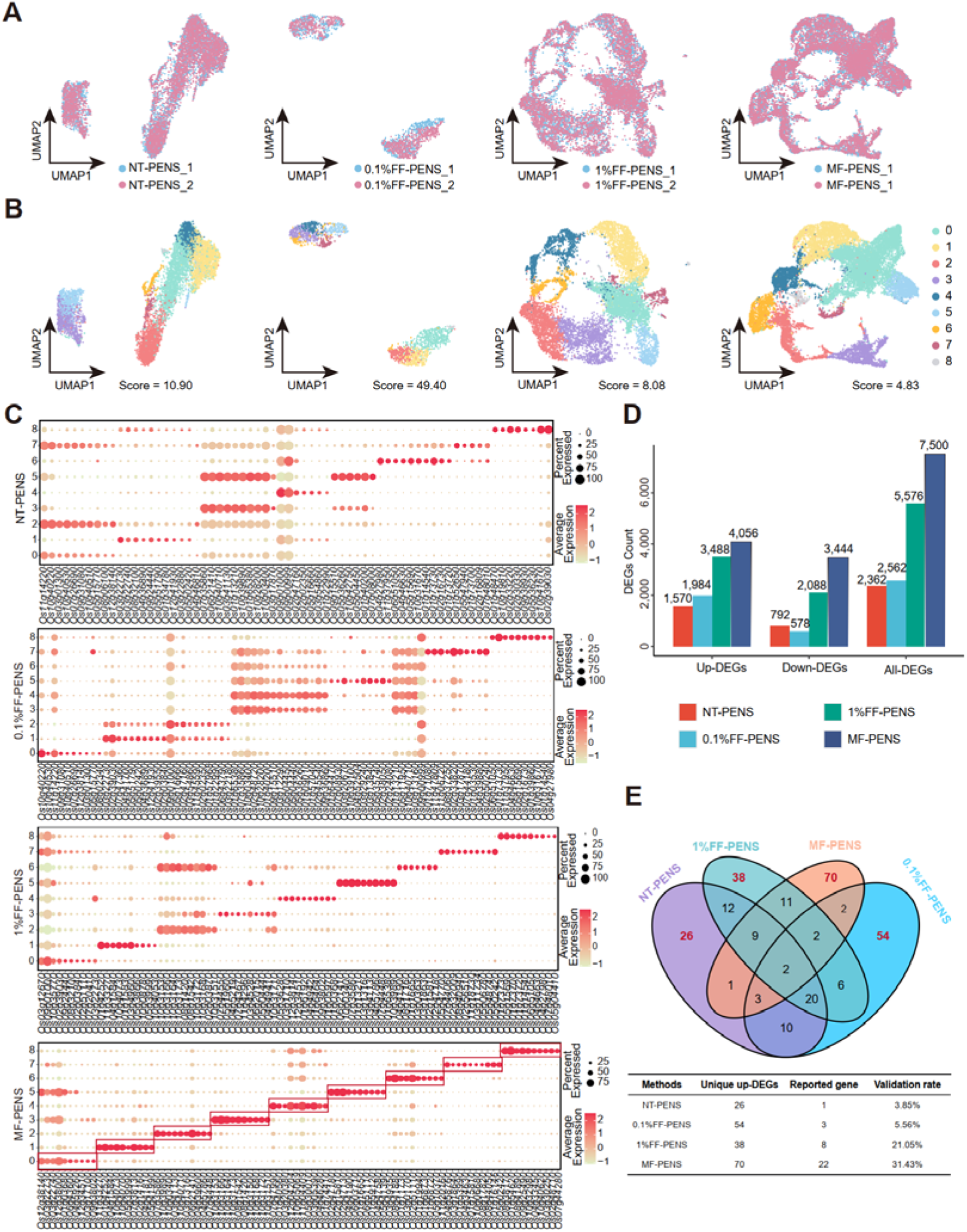
MF-PENS-generated snRNA-seq data exhibit superior cell clustering performance. **(A)** UMAP visualization of two biological replicates for each of the four methods (NT-PENS, 0.1%FF-PENS, 1%FF-PENS, and MF-PENS). **(B)** UMAP plots of Seurat clustering results, showing nine clusters derived from snRNA-seq data generated by the four methods, corresponding to **(A)**. Clustering performance was quantified using a transformed Modularity score, defined as -(Modularity x 10^5). Lower scores indicate superior clustering results. **(C)** Bubble plot of the top 10 upregulated differentially expressed genes (up-DEGs) in each cluster based on the clustering results across the four methods. **(D)** Bar plot of up-DEGs (*P_adj* < 0.05, log_2_FC > 0.25), down-DEGs (*P_adj* < 0.05, log_2_FC < -0.25), and total significant DEGs (All-DEGs, *P_adj* < 0.05, |log_2_FC| > 0.25) identified by the four methods. **(E)** Venn diagram showing the overlap of the top 100 up-DEGs among the four methods. Columns in table display the count of method-specific up-DEGs, the subset of these genes validated by the *PlantCellMarker* database, and the calculated validation rate for each method.

The advantages of MF-PENS in clustering resolution were further substantiated by functional enrichment analysis. Compared to other methods, MF-PENS captured a broader spectrum of biological pathways relevant to specific root cell types. For instance, in cluster 7, MF-PENS uniquely enriched for terms such as “response to external biotic stimulus” (GO:0043207), “intraspecies interaction between organisms” (GO:0051703), “response to bacteria” (GO:0009617)—functional categories consistent with the molecular signature of root epidermis cells involved in environmental sensing and defense.^40-42^ Similarly, cluster 0 derived from MF-PENS exhibited significant enrichment in meristem-associated pathways, including ribosome (ko03010), spliceosome (ko03040), and ribosome biogenesis in eukaryotes (ko03008), supporting the identification of this cluster as meristematic cells actively engaged in protein synthesis and transcriptional regulation **(Figures S1C-F, S2A-D)**.^43,44^ Collectively, these results demonstrate that the superior clustering fidelity of MF-PENS directly translates into more accurate and biologically meaningful cell type resolution.

We next performed differential expression analysis to evaluate marker specificity across the four methods. As visualized in the bubble plot of the top 10 upregulated DEGs (Up-DEGs) **(Figure 2C)**, DEGs detected from MF-PENS exhibited high specificity, with expression restricted to their corresponding cluster. Conversely, markers from the other three methods displayed diffuse expression patterns across multiple clusters. Statistically, MF-PENS significantly outperformed other methods in detecting upregulated, downregulated, and total differentially expressed genes **(Figure 2D)**. To assess biological fidelity, we analyzed the top 100 up-DEGs; MF-PENS identified the highest number of unique markers (70) compared to NT-PENS (26), 0.1%FF-PENS (54), and 1%FF-PENS (38) **(Figure 2E, Table S3-4)**. Crucially, cross-referencing these unique genes with the *PlantCellMarke*r database^45^ (https://www.tobaccodb.org/pcmdb/homePage) revealed that MF-PENS achieved the highest validation rate (31.43%) (**Table S2**), substantially surpassing other methods. These findings indicate that MF-PENS yields more biologically meaningful, cluster-specific signatures, enhancing downstream interpretability.

Collectively, these results highlight three pivotal advantages of MF-PENS in plant single-cell transcriptomics: (1) enhanced performance in cell clustering, (2) superior sensitivity in resolving cell-type-specific functional pathways, and (3) improved fidelity in identifying biologically validated DEGs. Consequently, MF-PENS not only ensures high data quality but also provides opportunity for more effectively mining deep biological insights into plant cellular heterogeneity.

### MF-PENS enables robust nuclei isolation from both fresh and frozen plant tissues

Compatibility with frozen samples is essential for experimental flexibility and the utilization of biobanked specimens. To validate the performance of MF-PENS on cryopreserved material, we selected rice roots and leaves as representative belowground and aboveground tissues. To mimic real-world transport and storage logistics, fresh samples were flash-frozen in liquid nitrogen, stored on dry ice for 15 days, and subsequently transferred to -80°C for another 15 days.

We rigorously benchmarked MF-PENS by processing matched fresh and frozen samples (n=2 biological replicates each) from root **(Figures 3A–G)** and leaf **(Figures 3H–P)** tissues, followed by snRNA-seq with stringent quality control.

**Figure 3.**
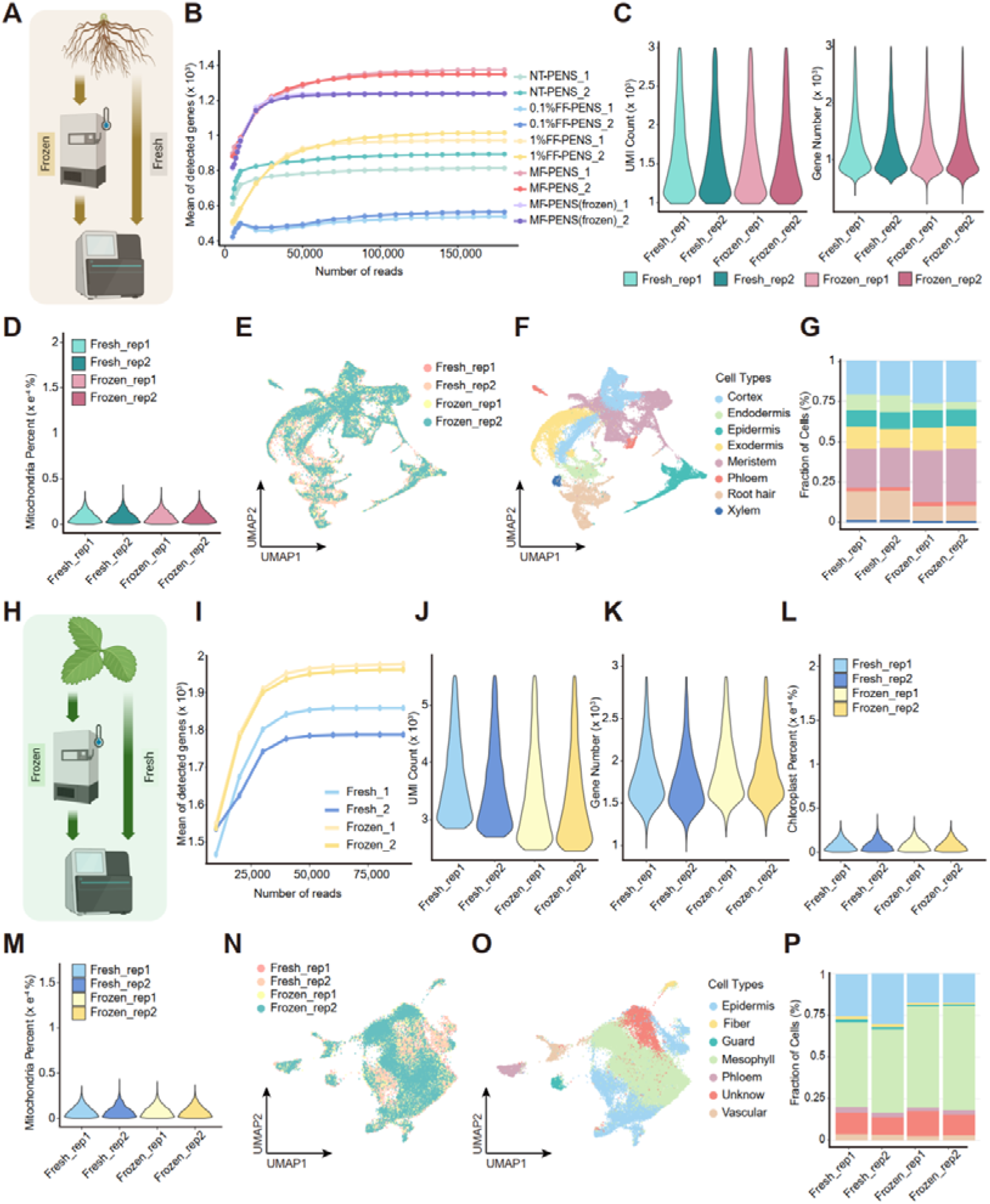
MF-PENS is applicable for both fresh and cryopreserved plant samples. **(A)** Workflow schematic of MF-PENS-based snRNA-seq for fresh and frozen rice root tissues. **(B)** Stepwise increases in mean detected genes and UMIs at different down-sampled sequencing depths for MF-PENS root data. **(C-D)** Violin plots showing UMIs, gene numbers, and mitochondrial percentages (y-axis × 10^-4^) for four samples, including two replicates each of fresh and frozen methods. **(E-F)** UMAP visualizations of integrated clustering and corresponding cell type annotation for four MF-PENS samples. **(G)** Proportion of cell types in **(F)** across four samples.**(H)** Workflow schematic of MF-PENS-based snRNA-seq for fresh and frozen rice leaf tissues. **(I)** Stepwise increases in mean detected genes and UMIs at different down-sampled sequencing depths for NT-PENS leaf data. **(J-M)** Violin plots showing UMIs, gene numbers, mitochondrial percentages (y-axis × 10^-4^%), and chloroplast percentages (y-axis × 10^-4^%) for four samples, including two replicates of fresh and frozen conditions. **(M-O)** UMAP visualizations of integrated clustering and corresponding cell type annotation for four NT-PENS samples. **(P)** Proportion of cell types in **(O)** across four samples.

In root tissue, MF-PENS achieved robust gene detection efficiency across all sequencing depths for both fresh and frozen replicates, confirming its suitability for transcriptomic profiling archival materials **(Figure 3B)**. Quantitative comparison of key quality metrics—including UMIs, detected genes, and mitochondrial gene percentages—showed no significant differences between fresh and frozen samples **(Figures 3C, D)**. These results demonstrate that MF-PENS effectively preserves transcriptomic integrity independent of storage conditions. Furthermore, UMAP confirmed the robust data integration, revealing distinct cell clusters with negligible batch effects attributable to storage conditions **(Figures 3E, S3A and S3B)**. Using reported canonical marker genes, we annotated eight distinct cell types **(Figures 3F, S4A-B)**. Cellular composition remained highly conserved between conditions, except for a specific reduction in root hair cells in frozen samples, likely due to physical shearing induced by cryopreservation **(Figures 3G)**.

Parallel experiments in leaf tissues yielded results concordant with those observed in roots. Fresh and frozen leaf samples exhibited high consistency across all key metrics, including the proportions of chloroplast-derived transcripts **(Figures 3I-M)**. Unsupervised clustering confirmed the effective integration of fresh and frozen samples, resolving 12 clusters that were successfully annotated to six major leaf cell types **(Figures 3N, O, S3C, S3D, S5A-B)**. Consistent with root tissues, cell type proportions in leaves remained largely stable across storage conditions **(Figure 3P)**.

Collectively, by consistently yielding high-quality snRNA-seq data from both fresh and frozen tissues (roots and leaves), MF-PENS demonstrates remarkable adaptability across diverse tissue types and storage conditions. This capacity to preserve transcriptomic integrity underscores the utility of MF-PENS for flexible experimental designs, enabling the effective exploitation of archived or transported plant resources.

### MF-PENS demonstrates broad versatility across diverse and recalcitrant plant tissues

Agronomically important crop tissues often harbor complex metabolic profiles.^46,47^ For instance, soybean seeds are replete with lipids^48^, whereas tomato fruits accumulate high concentrations of polysaccharides.^49,50^ Isolating high-quality nuclei from such recalcitrant tissues poses significant technical hurdles that severely impede snRNA-seq applications.^19,51^ To benchmark MF-PENS across diverse biochemical contexts, we selected four representative tissue types characterized by distinct physicochemical properties **(Figure 4A)**: infected tissue (soybean root nodules)^52,53^, oil-rich tissue (soybean seeds)^54^, polysaccharide-rich tissue (tomato fruit)^55^, and fragile tissue (maize callus).^56-58^. Following stringent quality control, MF-PENS consistently recovered high-quality nuclei from all biological replicates (n=2), yielding between 12,725 and 17,565 nuclei per library **(Figure 4B)**.

**Figure 4.**
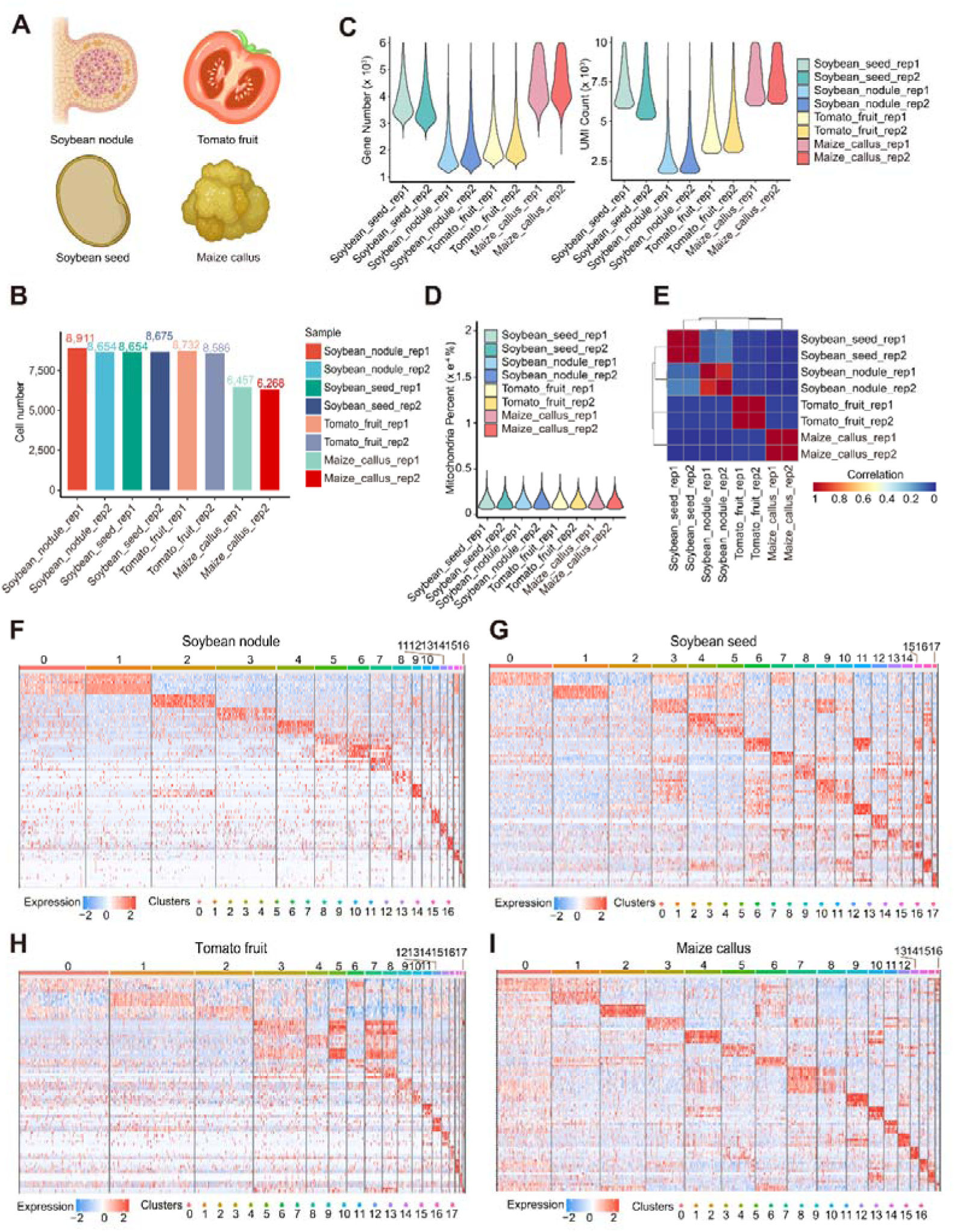
Universal applicability of MF-PENS across plant species and tissues. **(A)** Application of MF-PENS-based snRNA-seq to diverse tissue types, including infected tissue (soybean root nodules), polysaccharide-rich tissue (tomato fruit), lipid-rich tissue (soybean seeds), and callus tissue (maize callus). **(B)** Cell number of eight samples after quality control. **(C)** Violin plots of gene numbers and UMIs for eight biological samples (y-axis × 10^3^), including two replicates for each of the four tissue types. **(D)** Mitochondrial percentages (y-axis × 10^-4^%) for the eight biological samples. **(E)** Pearson correlation heatmap of the eight biological samples. **(F-G)** Heatmaps of the top five differentially expressed genes in each cluster identified from the four samples.

Sequencing metrics further confirmed the robust performance of MF-PENS across tissues of varying complexity. Specifically, soybean seeds yielded an average of 3,815 genes and 7,446 UMIs per nucleus. Soybean root nodules achieved an average of 2,136 genes and 3,128 UMIs per nucleus—a marked improvement over previous datasets, which reported averages of only ∼1,000–1,600 genes and ∼1,600–2,200 UMIs across developmental stages.^59,60^ Similarly, tomato fruits exhibited robust capture (2,349 genes; 5,137 UMIs), while maize callus demonstrated the highest efficiency, averaging 4,535 genes and 8,724 UMIs per nucleus. **(Figure 4C)**. Furthermore, mitochondrial gene proportions remained consistently low across all libraries **(Figure 4D)**, indicating minimal cytoplasmic contamination and efficient nucleus isolation. High concordance between biological replicates **(Figure 4E)** further corroborated the robustness of MF-PENS for profiling biochemically recalcitrant tissues.

To investigate tissue-specific gene expression patterns, we performed unsupervised clustering on the integrated datasets. UMAP visualization revealed distinct, well-separated cell populations **(Figures S3E-H)**, demonstrating the preservation of cell-type identities. Differential expression analysis identified robust cluster-specific gene expression signatures, visualized via heatmaps of the top up-DEGs **(Figures 4F-I)**. These potential markers exhibited exquisite cluster specificity with minimal background noise, highlighting the resolution of MF-PENS in dissecting transcriptional heterogeneity within complex tissues.

Collectively, these results demonstrate that MF-PENS effectively circumvents the technical barriers associated with metabolically complex plant materials. As a versatile and robust method, MF-PENS empowers researchers to generate high-quality single-cell data from recalcitrant tissues, thereby facilitating deeper insights into plant development and physiology.

### MF-PENS enables single-nucleus level dissection of plant heat stress responses

The scRNA-seq permits the high-resolution dissection of transcriptomic landscapes, unveiling lineage-specific activation patterns of key stress-response pathways.^61-63^ Beyond enhancing nuclear yield, MF-PENS confers a critical advantage via rapid metabolic inactivation, effectively "locks" the intracellular transcriptional state at the precise moment of sampling^64^. This instantaneous preservation mitigates transcriptional artifacts induced by mechanical stress or environmental fluctuations during processing, ensuring that sequencing data faithfully reflect the physiological status of the plant *in situ*. This capability enable MF-PENS serves as a robust, high-fidelity tool for elucidating the mechanisms of plant stress resilience. To evaluate this utility, we profiled rice leaves under normal (Control) and heat stress (Heat) conditions using both NT-PENS and MF-PENS, generating eight datasets across two biological replicates.

At the sample level, integrated analysis demonstrated successful batch correction and data integration for both methods **(Figures 5A, B)**. B Differential expression analysis (Control vs. Heat) revealed that MF-PENS captured a significantly broader repertoire of heat-responsive genes, with ∼80% of up-DEGs being exclusive to the MF-PENS datasets **(Figure 5C)**. Gene ontology enrichment (GO) analysis further confirmed that these 1,247 MF-PENS-exclusive genes were significantly enriched in “response to heat” (GO:0009408), whereas DEGs specific to NT-PENS lacked such functional relevance **(Figures 5D, E and Table S4)**. These findings underscore the superior sensitivity of MF-PENS in capturing the full spectrum of stress-responsive transcriptional changes.

**Figure 5.**
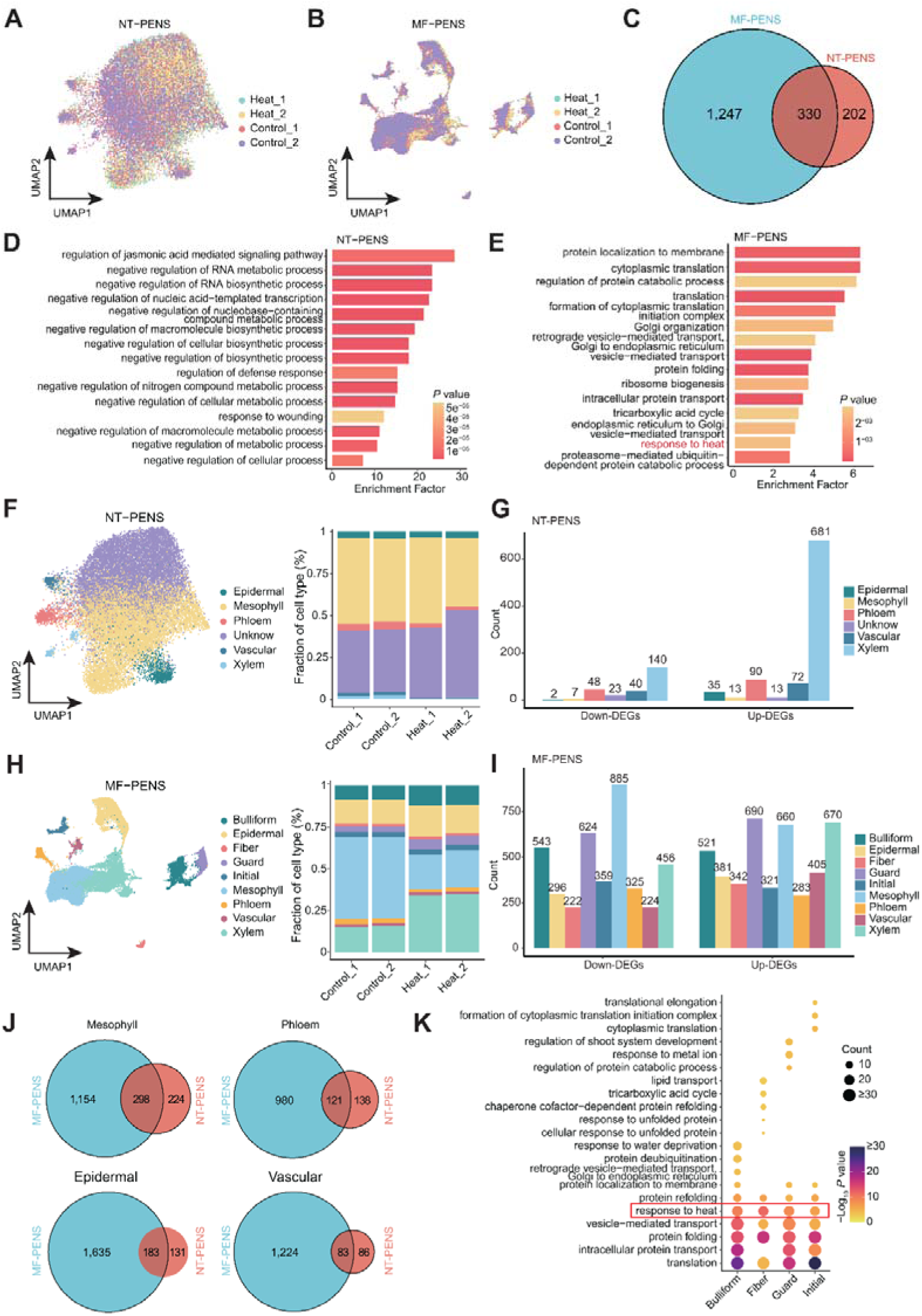
MF-PENS enables improved identification of cell types and detection of heat stress–responsive genes. **(A-B)** UMAP visualizations of integrated snRNA-seq data from NT-PENS and MF-PENS under control and heat stress conditions, each with two biological replicates. **(C)** Venn diagram showing the overlap of up-DEGs between control and heat stress conditions in NT-PENS and MF-PENS datasets. **(D-E)** Top 20 enriched biological processes based on the up-DEGs identified in **(C)**. **(F)** UMAP visualizations of cell type annotations and percentage based on clustering results from **(A)**. **(G)** Bar plot showing the number of up-/down-DEGs in each annotated cell type from **(F)**. **(H)** UMAP visualizations of cell type annotations and percentage based on clustering results from **(B)**. **(I)** Bar plot showing the number of up-/down-DEGs in each annotated cell type from **(H)**. **(J)** Venn diagram of heat-responsive genes (control vs. heat) shared by four commonly annotated cell types in both NT-PENS and MF-PENS datasets. **(K)** Bubble plot of the top 10 enriched biological processes of heat-responsive genes uniquely identified in four MF-PENS–specific cell types.

We next assessed resolution at the cell-type level via unsupervised clustering and canonical marker-based annotation. NT-PENS datasets annotated only five cell types and exhibited poor reproducibility between replicates, with minimal shifts in cellular composition observed under heat stress **(Figures 5F S8A)**. Furthermore, cell populations in NT-PENS appeared mixed on the UMAP, confounding downstream bioinformatic interrogation. In stark contrast, MF-PENS successfully delineated nine distinct cell lineages with high consistency across biological replicates**(Figures 5H, S8B)**.

At the level of transcriptional regulation, we computed DEGs for each annotated cell type for both methods **(Figures S6C, D)**. Under identical statistical thresholds, MF-PENS identified a substantially higher number of DEGs per cell type. This enhanced sensitivity suggests that MF-PENS captures cellular heterogeneity with greater fidelity, thereby accurately reconstructing cell-type-specific activation patterns **(Figures 5G, I)**. Comparative analysis of four shared cell types (mesophyll, phloem, epidermal, and vascular) revealed that MF-PENS-specific DEGs were significantly enriched in heat stress response-responsive pathways, while NT-PENS-specific DEGs showed no such functional enrichment **(Figures 5J, S8E-F, S7A-J and Table S5, 6)**. Additionally, DEGs identified within the four cell types uniquely resolved by MF-PENS (bulliform, fiber, guard, and initial cells) also exhibited robust enrichment for heat-response functions **(Figure 5K)**. These results further validate the superior capability of MF-PENS in resolving cell type-specific transcriptional regulation patterns under stress.

In summary, our multi-level analysis demonstrates that MF-PENS provides a comprehensive and high-fidelity representation of heat-induced transcriptomic remodeling. MF-PENS can serve as a robust, high-resolution platform for elucidating plant stress resilience mechanisms and environmental adaptation.

### MF-PENS enables precise cell subtyping and nuanced transcriptional profiles

Resolving cellular heterogeneity at the level of subpopulations is critical for elucidating lineage trajectories, functional compartmentalization, and the developmental roles of specific cell subsets within plant tissues.^19,65^ For instance, rice leaf epidermis comprises distinct upper and lower layers, each characterized by divergent gene regulatory networks and developmental functions.^66-68^ However, due to their high transcriptional similarity, previous scRNA-seq data struggled to reliably distinguish these subtypes.^69,70^ MF-PENS addresses this challenge by generating high-quality snRNA-seq data capable of robustly separating transcriptionally similar cell populations.

First, following the strategy described by Xia Keke *et al.*^66^ to distinguish upper and lower epidermal cells, we generated high-resolution Stereo-seq data from rice leaves. Based on cell wall morphology and spatial coordinates, we annotated upper and lower epidermal cells *in situ* as ground truth **(Figure 6A)**. We utilized these spatially resolved cells to derive specific up-DEGs for the upper and lower epidermis, establishing them as Stereo-seq-derived markers. Subsequently, we extracted all epidermal cells from NT-PENS (epidermal) and MF-PENS (bulliform, guard, and epidermal) datasets under normal growth conditions. Leveraging Stereo-seq-derived markers as highly variable gene (HVGs) for cell clustering, we successfully resolved two epidermal subpopulations for both datasets with highly consistent cell proportions across biological replicates **(Figures 6A-B, E-F)**. GO analysis revealed that cluster 0 in both datasets was significantly enriched for biological processes associated with upper epidermal cell function, including photosynthesis, light harvesting, and photorespiration **(Figures S8C, D, G)**, validating cluster 0 as upper epidermal cells. Using Stereo-seq-derived markers,^66^ we established the ground truth identities of the upper and lower epidermal cells for our snRNA-seq dataset. Accordingly, we used these validated epidermal cell identities as ground truth annotations for subsequent analysis **(Figures 6C, F)**.

**Figure 6.**
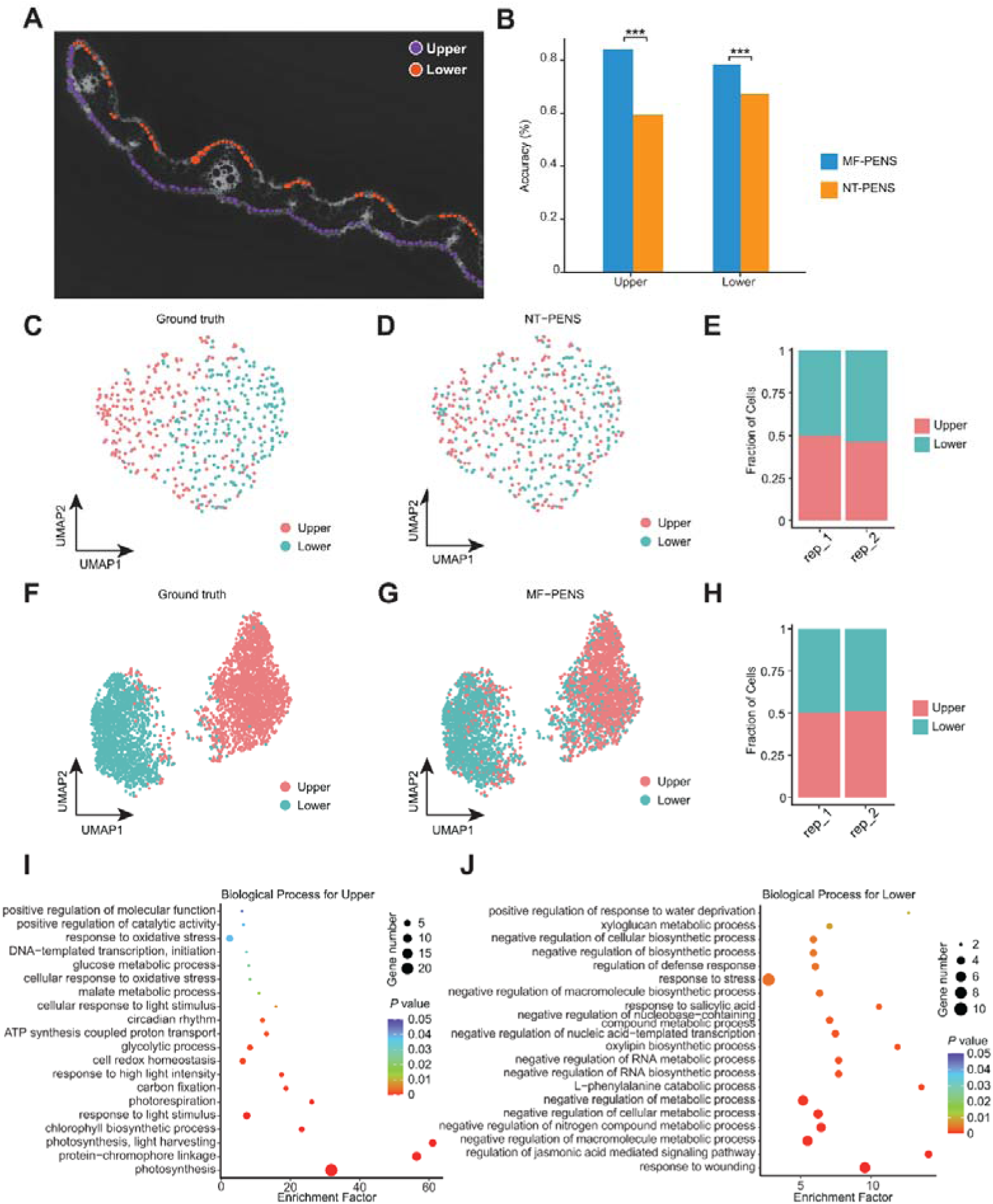
MF-PENS enables accurate identification of upper and lower epidermal cell subtypes in rice leaves. **(A)** Ground truth spatial locations of upper and lower epidermal cells in rice leaf tissue based on Stereo-seq. **(B)** Accuracy of epidermal cell subtype identification using MF-PENS and NT-PENS datasets. *** indicates a significant difference (*P* < 0.05). **(C)** Ground truth annotation of upper and lower epidermal cells in the NT-PENS dataset using a previously reported strategy. **(D)** Cell type annotation after standard clustering analysis by Seruat of the NT-PENS dataset. **(E)** Proportions of upper and lower epidermal cells across two biological replicates based on **(D)**. **(F-H)** Application of the same strategy to MF-PENS data: ground truth upper and lower epidermal cells **(F)**, clustering-based cell type annotation **(G)**, and cell type proportions across two replicates **(H)**. **(I-J)** Bubble plots showing the top 10 enriched biological processes for DEGs in the two epidermal subtypes identified in MF-PENS annotations **(G)**.

Next, we performed unsupervised clustering using Seurat,^71^ employing normal HVGs rather than Stereo-seq-derived markers on both datasets. The resulting clusters were then benchmarked against the Stereo-seq-defined ground truth labels. MF-PENS achieved a classification accuracy of 84% for lower epidermis and 78% for upper epidermis, significantly outperforming NT-PENS (59% and 67%, respectively) **(Figure 6B)**. UMAP visualization confirmed that MF-PENS clusters were discrete and aligned closely with ground truth labels, whereas NT-PENS data exhibited considerable mixing **(Figures 6D, G)**. This classification accuracy was highly reproducible across biological replicates, supporting the robustness of MF-PENS in preserving fine-grained transcriptional identities **(Figures 6E, H)**. Functional characterization of MF-PENS-derived clusters confirmed that the upper epidermal population was enriched for photosynthesis-related terms, while the lower epidermal population exhibited enrichment for biosynthetic and metabolic processes, consistent with their known functions^66^ **(Figures 6I, J, and Table S7-9)**.

In summary, MF-PENS effectively surmounts the resolution limits of conventional methods, enabling the high-precision identification of fine-grained cell subtypes. This capability establishes MF-PENS as a powerful tool for dissecting the functional heterogeneity and specialization of plant tissues.

## Discussion

Despite the transformative impact of single-cell technologies in animal research, their application in plant systems has been impeded by the structural rigidity of cell walls and the prevalence of interfering secondary metabolites. These physical and biochemical barriers often compromise nuclei integrity and yield, limiting the utility of scRNA-seq. To circumvent these hurdles, we developed MF-PENS, a novel nucleus isolation method that integrates methanol fixation with Percoll density gradient centrifugation. Unlike traditional protocols that rely on fresh tissue and FACS, our approach utilizes methanol fixation to rigidify the nuclear envelope and arrest transcriptional activity immediately upon sampling **(Figure 1A)**. MF-PENS demonstrates superior performance in preserving nuclear morphology and enhancing transcript capture efficiency compared to standard methods. Our data confirms that MF-PENS not only improves sequencing metrics—such as nuclear yield and integrity **(Figures 1B-E)**—but also significantly boosts the sensitivity of differential gene expression analysis **(Figure 2)**.

A critical limitation of current plant snRNA-seq protocols is the reliance on fresh tissue, which restricts experimental design and logistical flexibility. A pivotal advantage of MF-PENS is its compatibility with cryopreserved samples. We demonstrated that MF-PENS maintains high data fidelity in both fresh and frozen tissues across distinct organs (leaves and roots) **(Figure 3)**. This capability is transformative for plant science, as it enables the retrospective analysis of biobanked specimens and facilitates sample collection from remote field sites where immediate processing is impractical.

Furthermore, we validated the robustness of MF-PENS on biochemically complex, "recalcitrant" tissues—including polysaccharide-rich tomato fruits, lipid-laden soybean seeds, symbiont-containing root nodules, and polyphenol-rich maize callus **(Figure 4A)**. By successfully generating high-quality transcriptomic profiles from these diverse systems, MF-PENS proves its broad applicability, overcoming the metabolite interference that typically hampers plant nuclei isolation.

Beyond technical robustness, MF-PENS delivers the high-resolution fidelity necessary to dissect fine-scale biological heterogeneity. In rice leaves, the method achieved superior clustering consistency, resolving nine distinct cell lineages **(Figure 5H)**. Crucially, MF-PENS demonstrates pivotal utility in decoding plant stress responses: under heat stress, it exhibited exceptional sensitivity, capturing a comprehensive repertoire of heat-responsive genes and unveiling specific regulatory pathways that were obscured in lower-quality datasets **(Figures 5J-K)**. This capability is essential for elucidating the complex, cell-type-specific mechanisms underlying plant stress resilience.

Furthermore, we addressed the long-standing challenge of distinguishing transcriptionally similar cell subtypes, specifically the adaxial (upper) and abaxial (lower) epidermal layers. By leveraging Stereo-seq as referent ground truth, we benchmarked the resolving power of MF-PENS against NT-PENS. The results demonstrate that MF-PENS accurately discriminates between these closely related epidermal lineages, capturing distinct gene signatures and functional specializations **(Figure 6)**. This capacity to resolve subtle transcriptional differences underscores the method’s potential for investigating complex developmental lineages.

While MF-PENS consistently yields robust snRNA-seq data across a variety of tissues, several limitations remain. First, while we validated the method on four major crop species, its applicability to a wider range of plant taxa, particularly those with extreme secondary metabolite profiles, remains to be comprehensively assessed. Second, this study did not include samples with distinctive tissue characteristics such as bark, xylem, and strawberry fruit. Bark and xylem are highly lignified tissues with extremely thickened cell walls rich in lignin, cellulose, and other secondary metabolites, whereas tissues like strawberry fruit contain substantially larger nuclei. These features may impede efficient nuclear release, compromise nuclear integrity, or cause abnormal sedimentation during Percoll density gradient centrifugation. For such challenging tissues, we recommend targeted optimization strategies, specifically extending the fixation duration to 45–60 minutes and increasing the centrifugal force to 1000–1500 g during gradient centrifugation to ensure better structural preservation and recovery. In addition, for plant organs where obtaining sufficient sample material is challenging, we recommend homogenization using steel beads directly within the centrifuge tube. Compared to traditional mortar and pestle grinding, this method significantly reduces sample loss; in such cases, a minimal sample amount (just enough to cover the bottom of the tube, ∼0.1 g) is sufficient to meet the requirements for library construction. Third, it is important to note that the choice of fixative may introduce selection bias. Specifically, methanol fixation may affect nuclear recovery efficiency in certain cell types, such as highly vacuolated cells, potentially leading to their underrepresentation in the final dataset. Fourth, this study focused exclusively on transcriptomics. The compatibility of MF-PENS with other single-nucleus multi-omics modalities, such as snATAC-seq and snCUT&Tag, requires further experimental optimization and validation. Finally, our study evaluated only methanol and formaldehyde as fixatives and has not systematically compared or tested alternative fixatives—such as ethanol, acetone, or mixed fixative solutions—for their effects on nuclear integrity, RNA quality, and downstream applications, which represents an additional limitation of the current approach.

Notably, in the context of rapidly advancing spatial omics and multi-omics integration, MF-PENS demonstrates broad application potential. Of particular importance is its strong synergy with high-resolution spatial transcriptomics technologies such as Stereo-seq. While Stereo-seq provides precise spatial localization, its per-spot (e.g., at bin20 resolution, corresponding to a 10 µm² pixel) gene capture depth is generally lower than that of snRNA-seq. Our analyses show that the high-quality snRNA-seq data generated by MF-PENS exhibit strong concordance with Stereo-seq data in terms of cell type annotation and gene expression profiles **(Figure 6)**, effectively compensating for Stereo-seq’s limited transcriptomic coverage. This complementarity positions MF-PENS as an ideal upstream tool for integrated single-nucleus and spatial transcriptomics workflows, laying a solid foundation for future multidimensional characterization of cell types, functional states, and spatial organization in plant tissues.

## Conclusion

In summary, MF-PENS represents a low-barrier, high-performance solution for plant single-nucleus transcriptomics. By effectively addressing the challenges of nuclear isolation in metabolite-rich and frozen specimens, this method significantly expands the accessibility and scope of plant single-cell research. The enhanced sensitivity, reproducibility, and biological interpretability provided by MF-PENS pave the way for a paradigm shift from bulk tissue analyses to high-resolution, cell-type-specific investigations. We anticipate that the widespread adoption of MF-PENS will facilitate a deeper mechanistic understanding of plant development, stress resilience, and environmental adaptation, ultimately contributing to advancements in basic plant biology and sustainable crop improvement.

## Materials and methods

### Plant materials and growth conditions

The plant materials used in this study included: rice roots (*Oryza sativa* L. cv. Zhonghua 11, 3 days post-germination), rice leaves (*Oryza sativa* L. cv. Zhonghua 11, 14 days post-germination with an additional 3-hour treatment), soybean nodules (*Glycine max* L. cv. Williams 82, 24 days post-inoculation), soybean seeds *(Glycine max* L. cv. Zhonghuang 13, 1-day post-germination), tomato fruits (*Solanum lycopersicum* cv. Ailsa Craig, red fruits at 14 days post-anthesis), and maize callus tissues (*Zea mays* L. inbred line B73). For the salt-treated rice leaf samples, the culture medium was replaced with a solution containing 150 mM NaCl at 14 days post-germination, followed by an additional 3 hours of incubation before sample collection. The control group underwent the same procedure, except that a fresh culture medium without NaCl was used during the replacement step.

### Plant tissue fixation

In this study, a standard initial sample size of approximately 0.2 g (fresh weight) was used for all experiments.

#### No-treatment group

Plant samples were rapidly cut into ∼0.5 cm² fragments using a razor blade and subjected to no fixation. The tissue fragments were washed twice with NIBTA buffer (10 mM Tris-HCl (pH 8.0), 10 mM EDTA (pH 8.0), 80 mM KCl, 1 mM Spermine-4HCl, 1 mM Spermidine-3HCl, 2% PVP-40, 0.3% Triton X-100, 1 mM DTT, 1× ProtectRNA™ RNase Inhibitor, 1× Protease Inhibitor Cocktail) and immediately processed for subsequent nuclear isolation.

#### 0.1% formaldehyde fixation group

Plant samples were rapidly cut into ∼0.5 cm² fragments using a razor blade and fixed on ice for 30 minutes in ice-cold NIBTA buffer containing 0.1% formaldehyde. After fixation, the tissue was washed twice with ice-cold NIBTA buffer and immediately processed for nuclear isolation.

#### 1% formaldehyde fixation group

Plant samples were rapidly cut into ∼0.5 cm² fragments using a razor blade and fixed on ice for 30 minutes in ice-cold NIBTA buffer containing 1% formaldehyde. After fixation, the tissue was washed twice with ice-cold NIBTA buffer and immediately processed for nuclear isolation.

#### 100% methanol fixation group

Fresh plant samples were rapidly cut into ∼0.5 cm² fragments using a razor blade, or frozen plant samples were crushed in liquid nitrogen with a pestle into fragments no larger than 0.5 cm² (avoiding grinding into a powder), followed by fixation in ice-cold 100% methanol on ice for 30 minutes. After fixation, the tissue was washed twice with ice-cold NIBTA buffer and immediately processed for nuclear isolation.

### Nucleus isolation

The tissue was transferred to a petri dish on ice and chopped with a Gillette razor blade for approximately 2 minutes. The chopped tissue was resuspended in NIBTA buffer and homogenized on ice for 5 minutes. The homogenate was then filtered through a 40 μm strainer, and the filtrate was collected into a new 50 mL centrifuge tube. The tube was centrifuged at 1,260 g for 10 minutes at 4℃. The supernatant was discarded, and the tissue pellet was resuspended in 4 mL of NIBTA buffer. In a new 15 mL centrifuge tube, 70% Percoll solution (3.5 mL Percoll and 1.5 mL NIBTA buffer) was added, and the 4 mL resuspended tissue homogenate was carefully overlaid onto the 70% Percoll layer. The tube was centrifuged at 650 g for 30 minutes at 4℃, during which most nuclei banded at the interface between the NIBTA buffer and the Percoll layer. The nuclei band was gently collected into a new 15 mL centrifuge tube, 10 mL of NIBTA buffer was added, and the tube was centrifuged again at 1,260 g for 5 minutes. The nuclei pellet was washed twice with NIBTA buffer, then resuspended in PBS containing 0.04% BSA. After counting the nuclei, the suspension was kept on ice for later use (it is recommended not to exceed 30 minutes).

### Single-nucleus RNA library construction and sequencing

The single-nucleus RNA-seq libraries were prepared as previously described with DNBelab C Series High-throughput Single-Cell RNA Library Preparation Set V2.0 (MGI, 940-000508-00), and Sequencing was performed using DNBSEQ T7.

### Spatial transcriptomics library construction

Stereo-seq^72^ was used to construct spatial transcriptomics libraries. The leaf was embedded in OCT and sectioned into 10-micron slices. These slices were then adhered to Stereo-seq chips. The chips containing tissue sections were incubated at 37 °C for 3 min. Permeability, reverse transcription, tissue removal, cDNA release, and library preparation were next performed using the previous methods.^66^ Finally, DIPSEQ T10 was used for sequencing.

### Preprocessing of snRNA-seq data

The snRNA-seq data were processed and analyzed using DNBelab_C_Series_HT_single cell-analysis-software (version 2.0),^73^ an open-source and flexible pipeline. Sequencing reads were aligned to respective reference genomes: rice reads to MSU v7.0,^74^ soybean reads to GCF_000004515.6,^75^ tomato reads to GCF_036512215.1,^76^ and maize reads to GCF_902167145.1.^77^ Stringent quality control measures were implemented across all samples. Potential doublets or multiplets were identified and filtered using the DoubletFinder package,^78^ followed by the removal of unbound or apoptotic cells. We established strict inclusion criteria for cell quality: (1) cells expressing fewer than 200 or more than 6,000 genes were excluded; (2) cells with UMIs below 500 or exceeding 40,000 were removed; (3) cells demonstrating mitochondrial gene expression exceeding 5% of total expression were discarded. These rigorous quality control steps ensured the retention of high-quality cells for subsequent bioinformatics analysis.

### Cell clustering and visualization

We processed the snRNA-seq data using the Seurat v4.3.2 package.^71^ To normalize gene expression across cells, we applied the “NormalizeData” function, which utilizes a global-scaling normalization method. This approach involves dividing the gene expression values by the total expression per cell, scaling the results by a factor of 10,000 (default), and applying a natural log transformation to stabilize the variance. To capture the most biologically relevant transcriptional variations, we identified the top 2,000 highly variable genes using the “FindVariableFeatures” function. To ensure robust integration of datasets and minimize batch effects, we employed Canonical Correlation Analysis (CCA) through the “FindIntegrationAnchors” and “IntegrateData” functions, which align shared biological features across samples. For dimensionality reduction, we performed principal component analysis (PCA) using the “RunPCA” function (npcs = 40) to extract the most informative components of variation. Subsequently, we applied graph-based clustering with the Louvain algorithm via the “FindClusters” function (resolution = 0.3) to group cells based on their transcriptional profiles. Finally, to visualize the high-dimensional data in an interpretable two-dimensional space, we utilized Uniform Manifold Approximation and Projection (UMAP) with the “RunUMAP” function (dims = 1:40, min.dist = 0.3). This approach preserves the local and global structure of the data, enabling clear visualization of distinct cell clusters and their relationships.

### Cell type annotation of rice leaf and root

In the rice leaf datasets, both NT-PENS and MF-PENS datasets were clustered into ten clusters **(Figure S3B)**, which were subsequently assigned to specific cell types. Epidermal cells were identified in clusters 3 and 4 based on the marker genes *Os03g52390* and *Os02g39930*.^79^ Cluster 8 was annotated as fiber, supported by the specific expression of the fiber marker genes *Os11g10310*, *Os11g10320*, *Os12g02320*, and *Os11g37900*.^70^ Cluster 9 was assigned as guard cell, as indicated by the high expression of the guard cell marker *Os03g10090*.^70^ Cluster 7 was annotated as phloem, characterized by the expression of *Os09g28510*, *Os05g03934*, and *Os04g14220*.^70,79^ Cluster 6 was identified as vascular, based on the expression of *Os10g14870*.^70^ Finally, clusters 0 and 1 were annotated as mesophyll, supported by the expression of *Os12g17600*, *Os01g41710*, and *Os07g37240*.^80^

In the rice root datasets, NT-PENS and MF-PENS were also clustered into ten clusters **(Figure S3D)**, and cell types were similarly annotated. Cluster 7 was assigned as phloem, supported by the marker genes *Os09g28510*, *Os05g34980*, *Os05g04000*, and *Os02g37070*.^45^ Cluster 0 was annotated as cortex, based on the expression of *Os11g02350*, *Os09g12970*, *Os04g59260*, *Os01g68580*, *Os12g02300*, *Os04g46810*, *Os02g44310*, and *Os01g68589*.^45^ Clusters 6 and 8 were identified as endodermis, characterized by three marker genes including *Os07g35480*, *Os06g13560*, and *Os05g12630*.^45^ Cluster 9 was annotated as xylem, supported by the expression of nine xylem-specific markers such as *Os01g62490*, *Os01g54620*, and *Os01g06580*.^45^ Cluster 3 was assigned as exodermis, based on the Plant Single Cell Transcriptome Hub (PsctH: http://jinlab.hzau.edu.cn/PsctH/) markers *Os06g29994*, *Os04g41740*, and *Os03g02460*.^81^ Cluster 5 was annotated as epidermis, supported by the expression of *Os10g31510*, *Os10g31660*, *Os03g06040*, and *Os06g20150*.^79^ Cluster 4 was identified as meristem, marked by *Os01g61920* and *Os03g02780*.^79^ Finally, cluster 2 was assigned as root hair, supported by nine specific markers including *Os10g42750*, *Os07g44460*, and *Os04g57860*.^79,81^

### Identification and enrichment analyses of DEGs

To identify differentially expressed genes (DEGs) across different treatments, we utilized the “FindMarkers” and “FindAllMarkers” functions in Seurat. Significant DEGs for clusters and each cell type were identified using a threshold of *P_*adjust < 0.05 and |log_2_ fold change| > = 0.25. In addition, Gene Ontology (GO) and Kyoto Encyclopedia of Genes and Genomes (KEGG) pathway analyses were performed on the DEGs to uncover their potential functional enrichments and biological signal transductions. The former includes biological processes, cellular components, and molecular functions, while the latter includes metabolism, environmental information processing, organismal systems, genetic information processing, and cellular processes. Functional enrichment analyses were conducted using the clusterProfiler R package,^82^ with significantly enriched terms identified based on a false discovery rate (FDR) threshold of < 0.05.

### Analysis of rice leaf Stereo-Seq data

In the Stereo-Seq data of rice leaves, we employed Calcofluor White (CW) staining images to visualize the cell walls and ssDNA staining images to reveal nuclear distribution, thereby capturing the structural features of plant tissue sections more comprehensively. Using CW-stained images aligned with the sequencing data, we delineated the boundaries of individual cells and assigned cell type labels, thus constructing ground-truth spatial transcriptomic data. To accurately identify and trace the complete cell wall contours of each epidermal cell, we combined manual cell outlining with the Cellpose^83^ automated cell segmentation software. This approach enabled precise characterization of the boundaries of both upper and lower epidermal cells in rice leaves **(Figure 6A)**. The spatial position of each cell on the sequencing chip was represented by its centroid coordinates. These cell data served as the ground truth for evaluating the ability of MF-PENS to discriminate between different epidermal cell types.

### Differential Expression Analysis and Benchmarking of Epidermal Subpopulations

We first performed differential expression analysis on spatially resolved cells identified via Stereo-seq. Using Scanpy, we conducted a Wilcoxon rank-sum test to identify significantly up-DEGs specific to the upper and lower epidermal layers, respectively (*P_adj* < 0.05, |log2FC| > 0.25). These Stereo-seq-derived up-DEGs were subsequently employed to establish ground truth references for the snRNA-seq datasets. We extracted all epidermal cells from the NT-PENS dataset (annotated as epidermal cells) and the MF-PENS dataset (annotated as bulliform, guard, and epidermal cells). Instead of unsupervised feature selection, we manually set the identified Stereo-seq up-DEGs as the “VariableFeatures” for these subsets. The data were then processed using the standard Seurat pipeline: normalization and scaling via “ScaleData”, dimensionality reduction via “RunPCA” and “RunUMAP”, neighbor identification via “FindNeighbors”, and clustering via “FindClusters”. The clustering resolution was adjusted to force the separation into two distinct clusters, serving as the ground truth reference for upper and lower epidermal identities **(Figures 6C and 6F)**. In parallel, we performed standard unsupervised clustering on the same snRNA-seq subsets. For this analysis, variable features were identified computationally using the “FindVariableFeatures” function in Seurat (parameters: “selection.method” = ‘dispersion’, “nfeatures” = 800). The subsequent workflow (scaling, PCA, UMAP, and clustering) remained identical, yielding clustering results derived solely from the snRNA-seq data structure **(Figures 6D and 6G)**. By comparing the unsupervised snRNA-seq clustering results against this Stereo-seq-derived ground truth, we evaluated the effectiveness of MF-PENS in resolving epidermal cell subtypes.

### Quantitative and statistical analysis

All experiments were repeated at least two times, and the data are presented as the mean ± standard deviation (SD). The significance of group differences was evaluated with a two-tailed Student’s t test or analysis of variance.

## Data availability

All datasets generated in this study have been submitted to the China National GeneBank (CNGB, accession number CNP0007006). All the other data and materials are available from the corresponding authors upon request.

## Code availability

We utilized the analysis of snRNA-seq data can be obtained from the corresponding author upon request. We also thank the China National GenBank (CNGB) for their generous support.

## Acknowledgments

We thank X. Lin (Institute of Genetics and Developmental Biology, Chinese Academy of Sciences) for providing the B73, Williams 82, and Zhonghuang 13 materials; F. Zhang (Institute of Genetics and Developmental Biology, Chinese Academy of Sciences) for providing the Zhonghua 11 materials; and T. Lin (College of Horticulture, China Agricultural University) for providing the Alisa Craig materials. This work was supported by the Biological Breeding-National Science and Technology Major Project [No. 2023ZD04076], the High-quality Science and Technology Journal Construction Project of Guangdong Province [No. 2025B1212100003] and [No. 2025B1212070003].

## Author contributions

J.-M.K., H.Y. and Z.L. designed the research. J.-M.K. and L.D. wrote the manuscript. J.-M.K., Q.L., and L.D. developed and tested the MF-PENS technology. J.-M.K., Q.L., M.-Y.Z., C.T. and M.-J.Z. did experiments. L.D., J.-M.K. and H.-L.Z. performed data analysis. Q.L., H.-L.Z., F.-Y.L., H.Y. and Z.L. revised the manuscript. L.D. and J.-M.K. prepared all the figures. All authors discussed the results and commented on the manuscript.

## Supplementary figures

**Figure S1.**
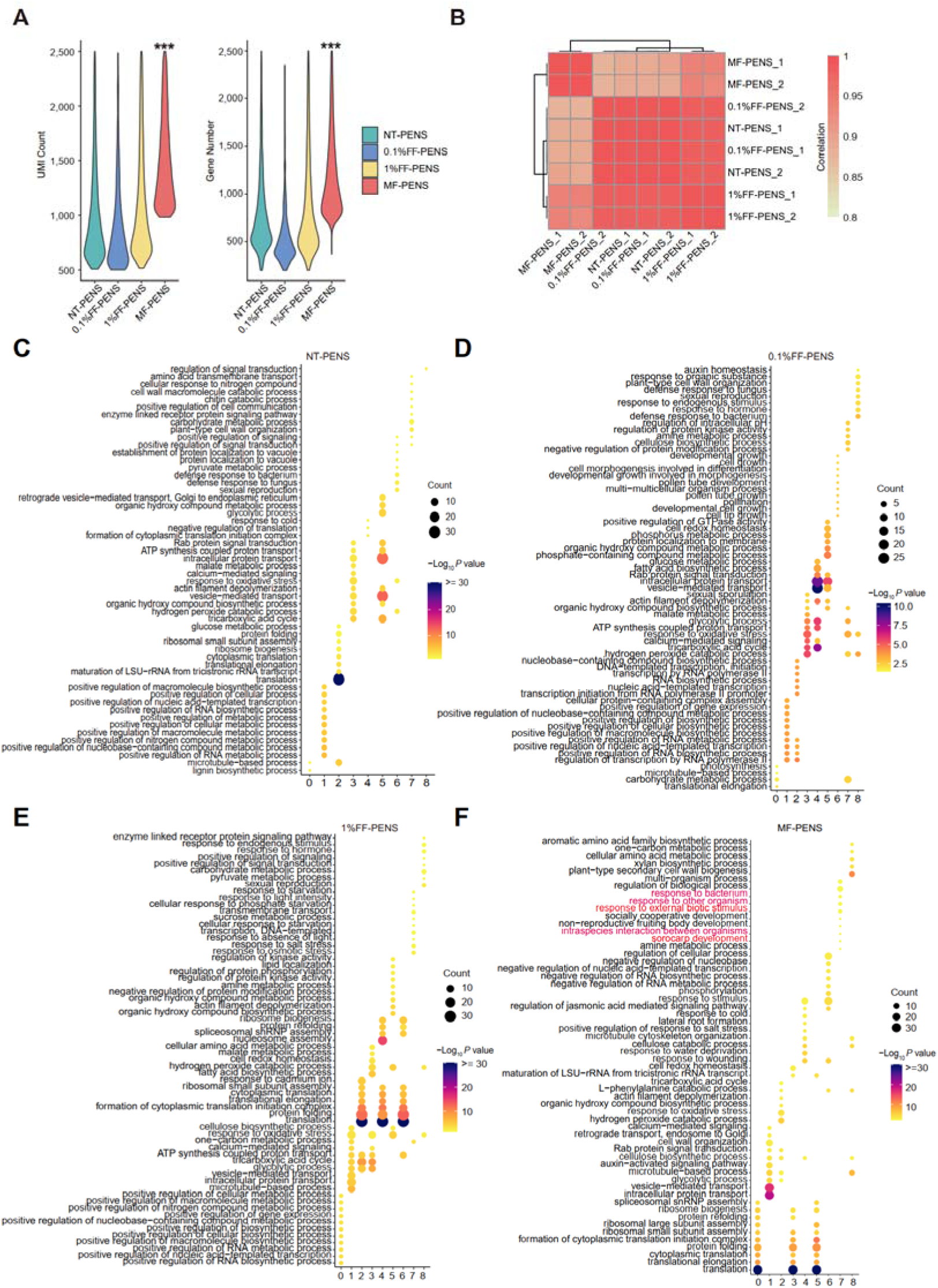
MF-PENS is the optimal method for single-nucleus sequencing in root tissue samples. **(A)** Violin plots showing UMIs and gene numbers across four methods. **(B)** Pearson correlation heatmap of eight samples where color gradients from blue to red indicate correlation coefficients from 0.5 to 1.0. Hierarchical clustering was applied to the correlation matrix. **(C-F)** Bubble plots showing the top 10 enriched biological processes of up-DEGs in each of the nine cell clusters identified from rice root using four different methods. The significance scale (-log_10_(*P* value) was capped at a maximum value of 30 (values >30 are displayed as 30). **Related to Figure 1 and Figure 2**.

**Figure S2.**
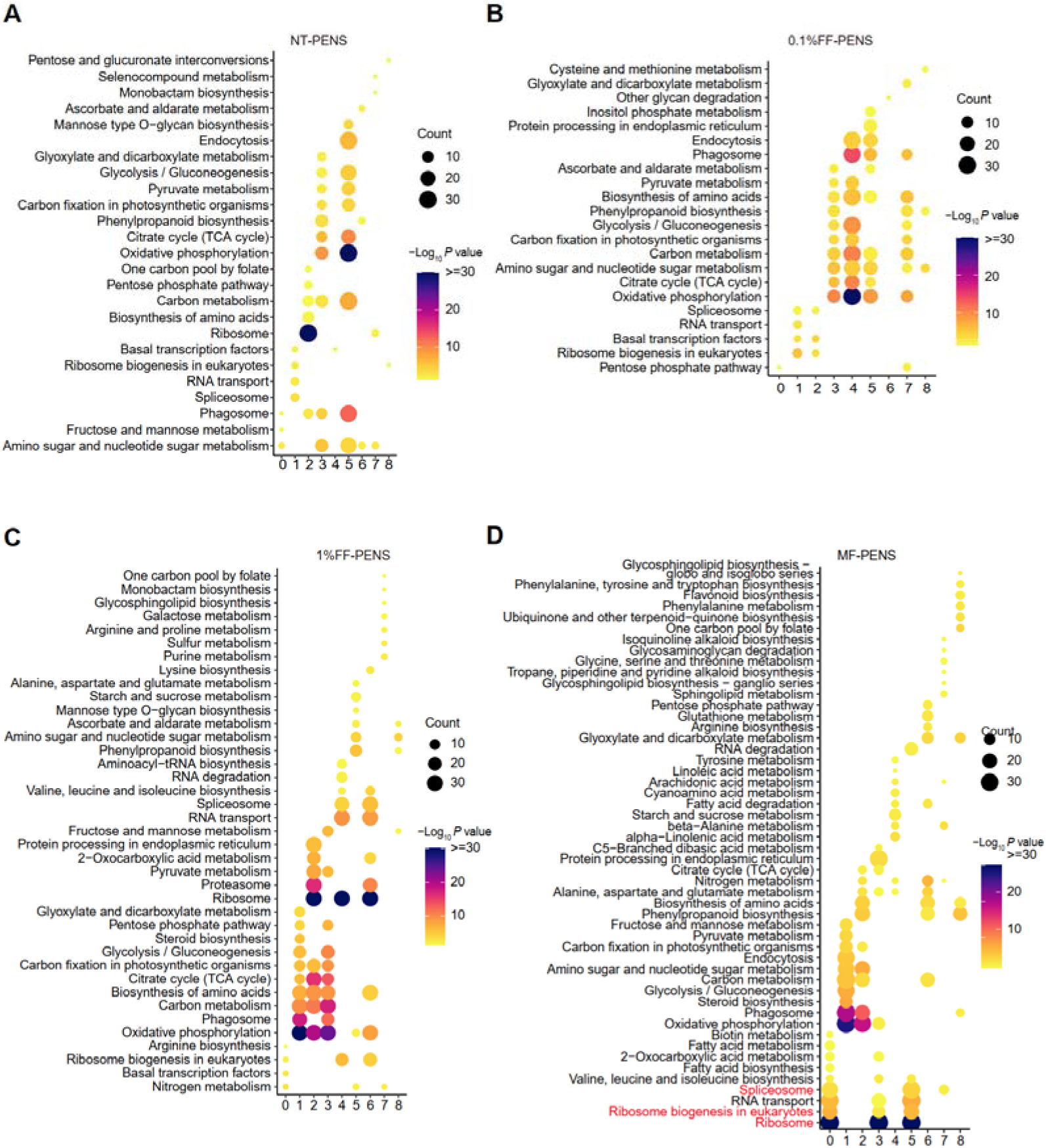
Comparative KEGG pathway enrichment analysis of rice root cell clusters across four nuclear isolation methods. Bubble plots display the top 10 significantly enriched KEGG pathways for the up-DEGs in each of the nine cell clusters identified by NT-PENS, 0.1%FF-PENS, 1%FF-PENS, and MF-PENS, respectively. The size of the dots typically represents the gene count or ratio, while the color gradient indicates the statistical significance level. For visualization clarity, the significance scale (-log_10_(*P* value) was capped at a maximum value of 30 (values >30 are displayed as 30). **Related to Figure 1 and Figure 2**.

**Figure S3.**
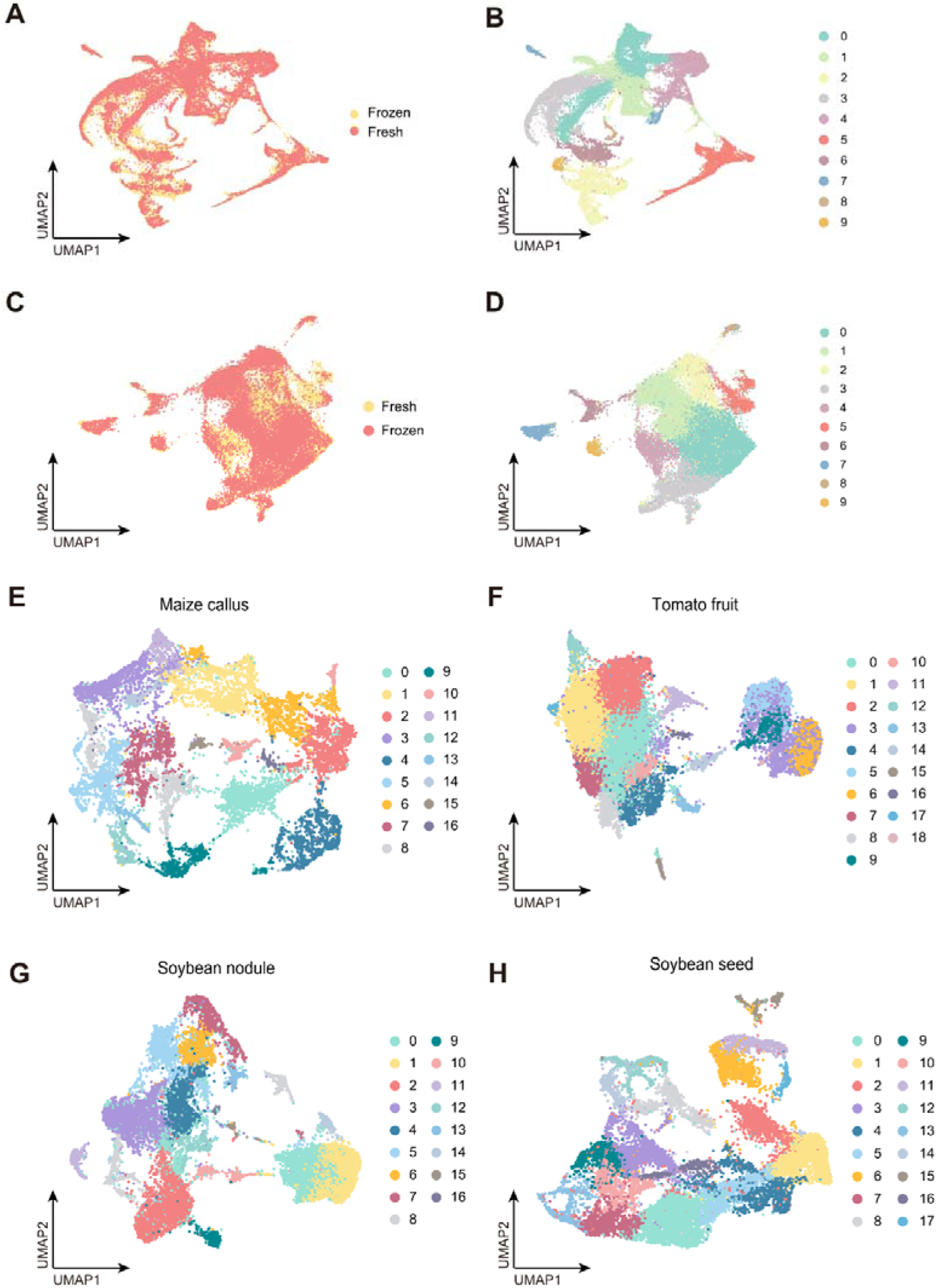
Cell clustering analysis of MF-PENS snRNA-seq data from diverse plant tissues. **(A-B)** UMAP visualization of integrated fresh and frozen rice root samples using CCA and the resulting cell clustering. **(C-D)** UMAP visualization of integrated fresh and frozen rice leaf samples using CCA and the resulting cell clustering. **(E-H)** UMAP plots showing cell clustering of snRNA-seq data from four challenging tissue samples. **Related to Figure 3 and Figure 4**.

**Figure S4.**
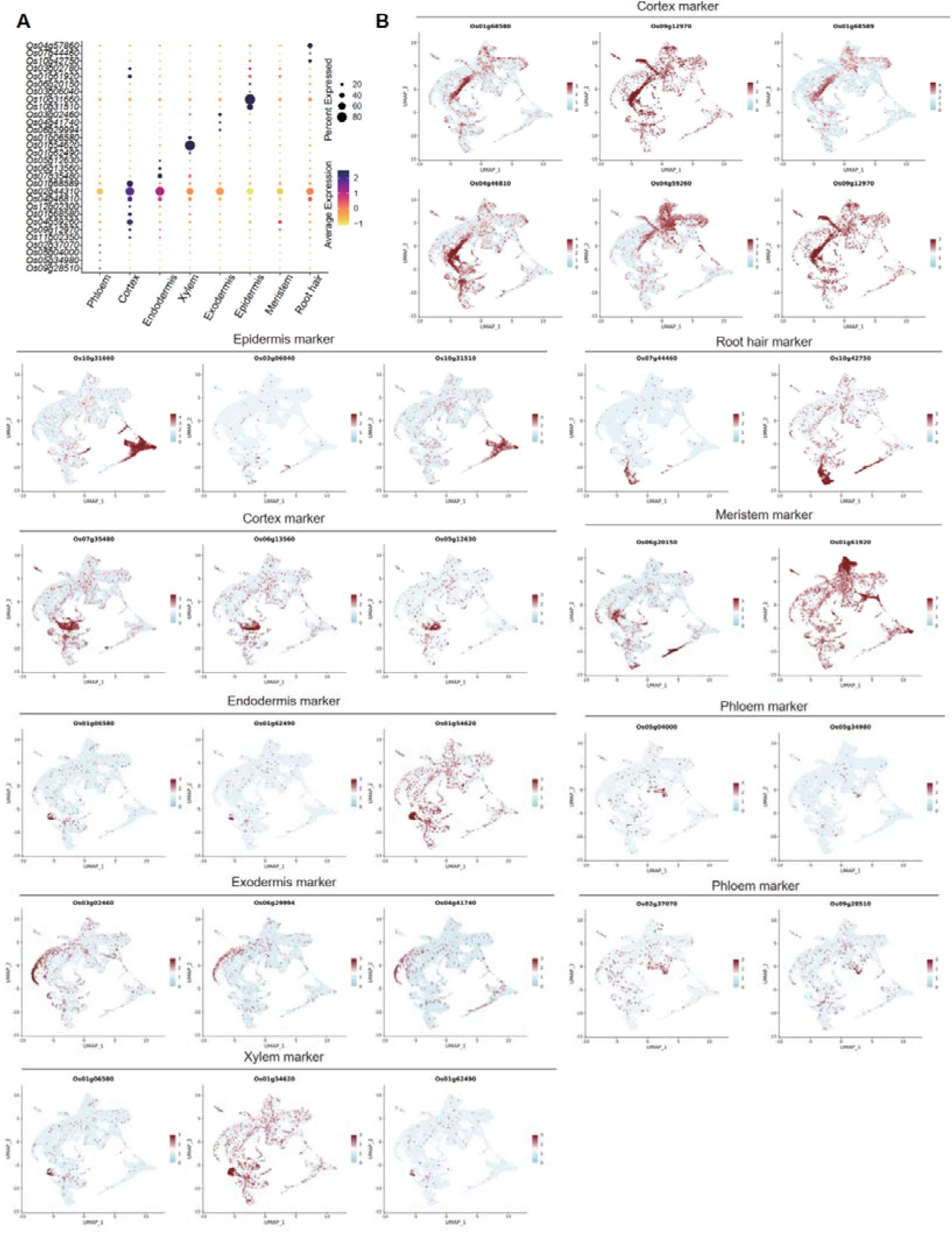
Expression profiles of marker genes used for cell type annotation in rice root snRNA-seq data. **(A)** Dot plot illustrating the expression patterns of representative marker genes across eight identified cell types. **(B)** Feature plots displaying the distribution and expression levels of 29 selected marker genes. The color gradient indicates the relative expression level. **Related to Figure 3**.

**Figure S5.**
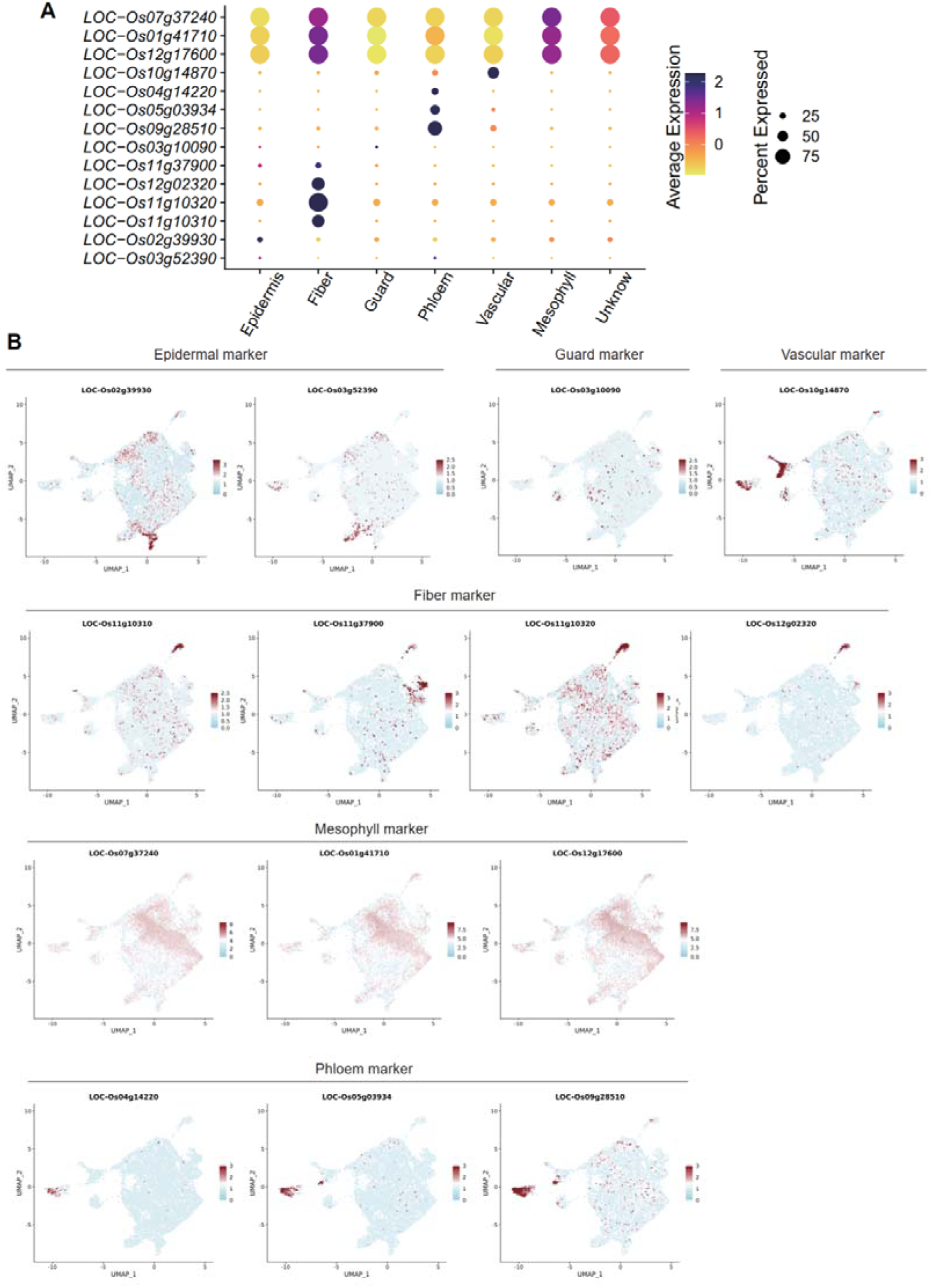
Expression profiles of marker genes used for cell type annotation in rice leaf snRNA-seq data. **(A)** Dot plot illustrating the expression patterns of representative marker genes across six identified cell types. **(B)** Feature plots displaying the distribution and expression levels of 14 selected marker genes. The color gradient indicates the relative expression level. **Related to Figure 3**.

**Figure S6.**
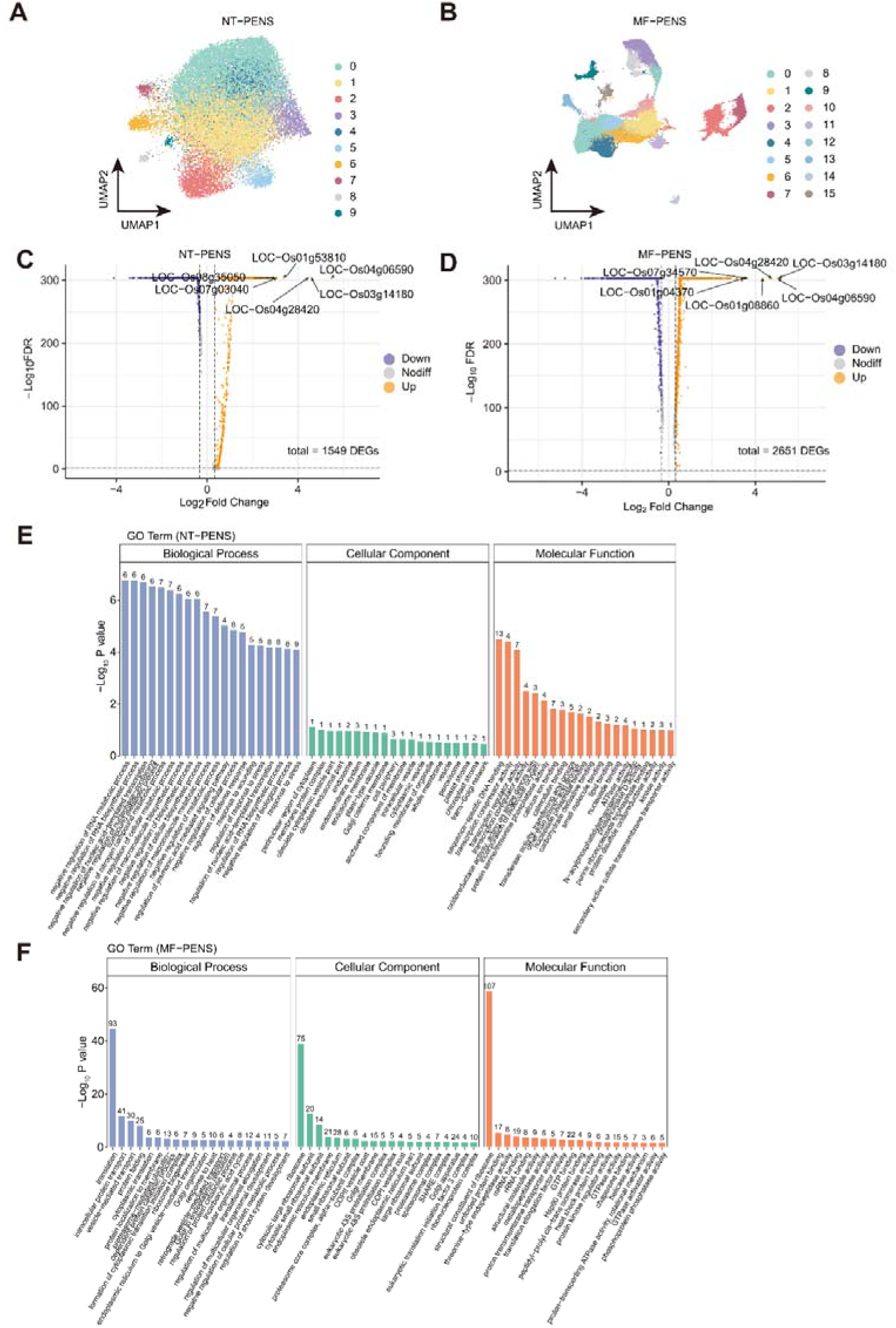
MF-PENS outperforms NT-PENS in capturing transcriptomic responses to heat stress in rice leaves. **(A–B)** UMAP plots of cell clustering of epidermal cells in rice leaves profiled by NT-PENS **(A)** and MF-PENS **(B)**. **(C-D)** Volcano plots of differentially expressed genes identified within each cluster from NT-PENS **(C)** and MF-PENS **(D)**, corresponding to **(A)** and **(B)**, respectively. **(E-F)** Top 20 enriched Gene Ontology (GO) terms derived from heat-responsive genes identified in Figure 5A-B, including categories of Biological Process (BP), Cellular Component (CC), and Molecular Function (MF). **Related to Figure 5**.

**Figure S7.**
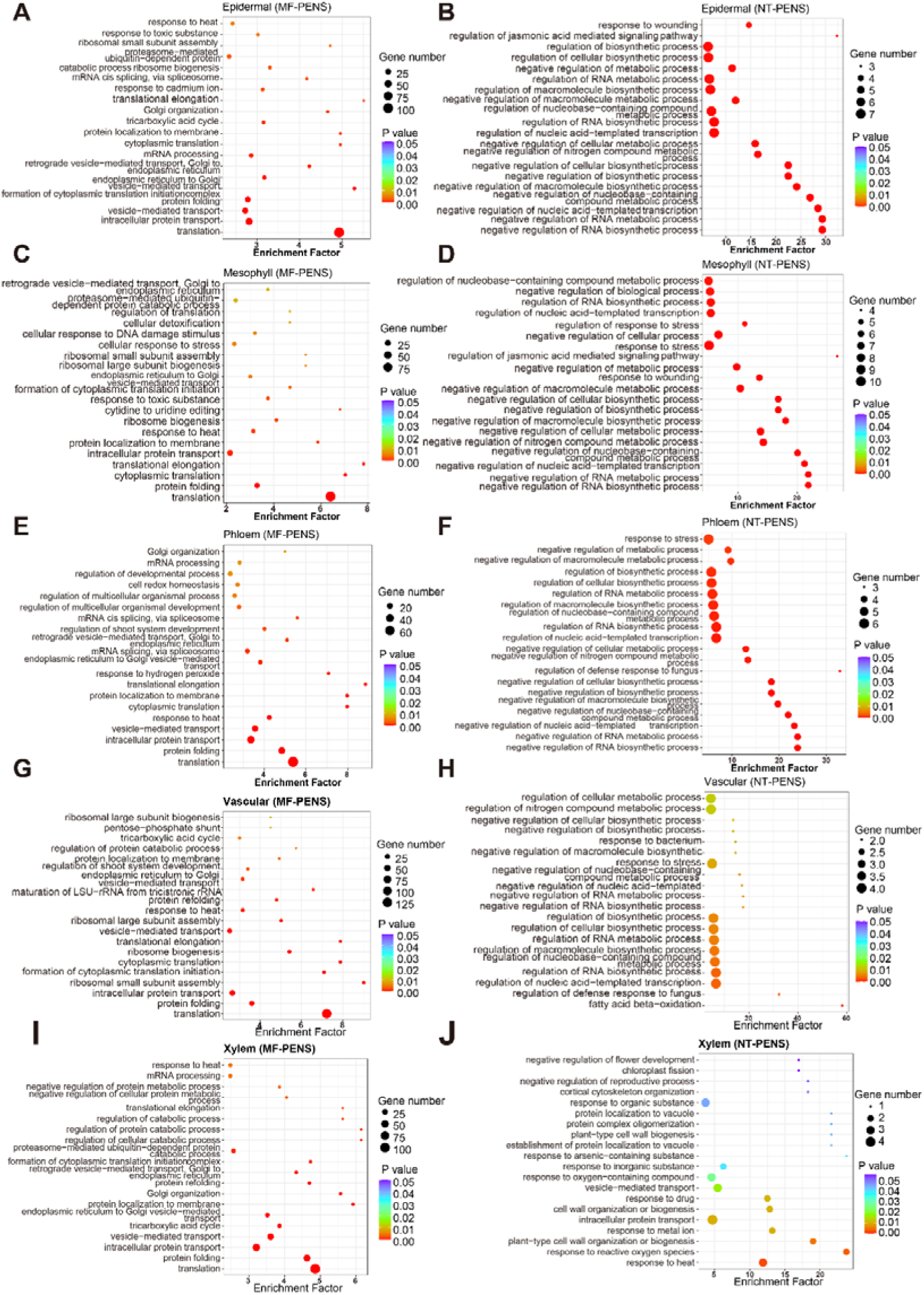
Bubble plots of the top 20 enriched biological processes for heat stress-responsive genes in rice leaf snRNA-seq data. The analysis was performed on five cell types shared between NT-PENS and MF-PENS datasets, comparing control versus heat stress conditions. **Related to Figure 5**.

**Figure S8.**
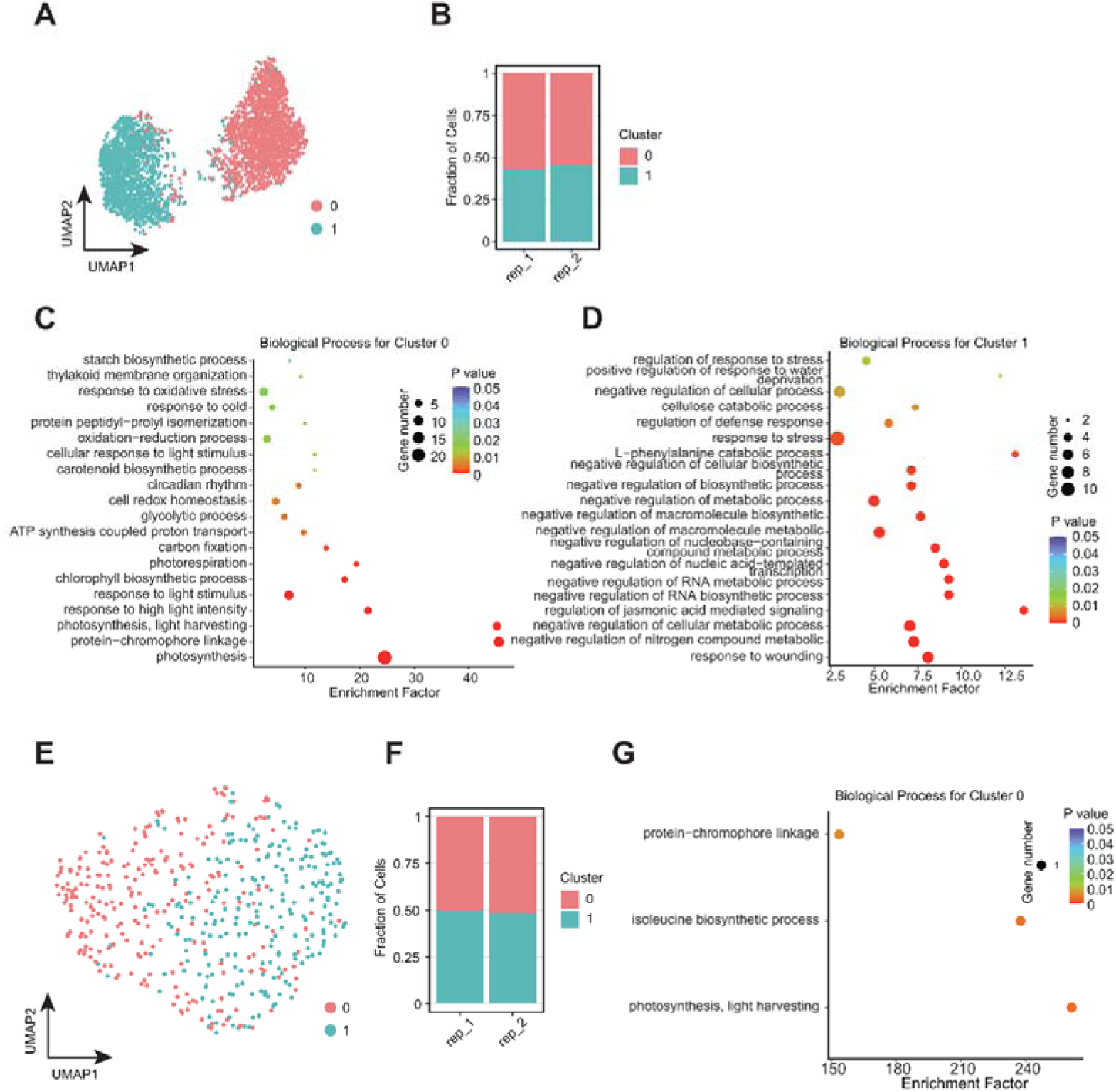
Identification of upper and lower epidermal cells in rice leaves using Stereo-seq data as ground truth. **(A)** UMAP plot of cell clustering from MF-PENS data, using marker genes of upper and lower epidermal cells derived from Stereo-seq as feature genes. **(B)** Proportions of the identified clusters across two biological replicates in MF-PENS data. **(C-D)** GO enrichment analysis of cluster-specific genes for the two clusters identified in **(A). (E)** UMAP plot of cell clustering from NT-PENS data using the same Stereo-seq-derived marker genes as features. **(F)** Proportions of clusters across two biological replicates in NT-PENS data. **(G)** GO enrichment analysis of cluster-specific genes from cluster 0 in **(E)**; no cluster-specific genes were identified for cluster 1. **Related to Figure 6**.

